# Evolutionary responses and genomic consequences of polyploidization in natural populations of *Orychophragmus*

**DOI:** 10.1101/2025.03.24.644964

**Authors:** Qiang Lai, Zeng Wang, Changfu Jia, Xiner Qumu, Rui Wang, Zhipeng Zhao, Yao Liu, Yukang Hou, Jianquan Liu, Pär K. Ingvarsson, Jing Wang

**Affiliations:** Key Laboratory for Bio-Resources and Eco-Environment of Ministry of Education, College of Life Sciences, Sichuan University, Chengdu, China; Linnean Centre for Plant Biology, Department of Plant Biology, Uppsala BioCenter, Swedish University of Agricultural Sciences, Uppsala, Sweden

**Keywords:** demographic history, gene expression, genetic load, *Orychophragmus*, polyploidization, selection efficacy

## Abstract

Polyploidization has occurred throughout the tree of life and is particularly common in plants. Despite its ubiquity, our understanding of the short- and long-term effects and consequences of genome doubling in natural populations remains incomplete. In this study, we identified a novel ploidy-variable species system within the ornamental and industrial oilseed genus *Orychophragmus* (Brassicaceae), which comprises six species, including diploid and tetraploid cytotypes of *O. taibaiensis*. By integrating population-scale genomic and transcriptomic datasets across the species in this genus, we constructed a robust phylogenetic framework and investigated the divergence and demographic history of *O. taibaiensis* in comparison to its relatives. Specifically, we characterized the geographical distribution patterns of diploids and tetraploids in natural populations of *O. taibaiensis*, confirmed the autopolyploid origin of tetraploids, and inferred their origin time relative to diploid counterparts. Our findings further revealed that, following genome doubling, tetraploids accumulated a higher genetic load of deleterious mutations, likely due to relaxed purifying selection facilitated by allelic redundancy. Additionally, genome doubling was associated with pronounced changes in gene expression patterns, with differentially expressed genes evolving under relaxed selective constraints. These results highlight that the initial masking of deleterious mutations, changes in expression regulation, and divergent efficacy of selection likely all contribute to shaping the establishment and evolutionary potential of polyploids.

## Introduction

Polyploidization, resulting from whole-genome duplication (WGD), has long been regarded as a key driver of plant speciation, adaptation to novel and extreme environments, and genetic innovation (Otto and Whitton 2000; Van de Peer, et al. 2017). While allopolyploidy has received considerable attention, autopolyploidy remains markedly less studied, despite its prevalence and suitability for exploring the direct impacts of immediate genome doubling without the complications associated with hybridity (Parisod, et al. 2010; Barker, et al. 2016). Autopolyploidy has been repeatedly shown to arise and establish from diploid populations, often leading to the coexistence of tetraploid and diploid populations within a single species (Monnahan, et al. 2019; Morgan, et al. 2020). However, our understanding of the evolutionary dynamics and genomic consequences of genome doubling in these populations remains limited (Kolář, et al. 2017; Mortier, et al. 2024). For example, it is unclear to what extent the additional chromosomal copies resulting from WGD can mask deleterious mutations, thereby influencing the genetic load (Griswold 2021). Furthermore, the degree to which genome duplication alters the efficiency of selection processes in tetraploid populations compared to their diploid progenitors remains largely unexplored (Spoelhof, et al. 2017). Polyploidy is also frequently associated with morphological and ecological changes, but the extent to which autopolyploidy reshapes gene expression patterns and the functional roles of differentially expressed genes remains an open and intriguing question (Osborn, et al. 2003).

Species diversification within the Brassicaceae family is intricately linked to repeated cycles of WGDs, with polyploidy being a common feature across many lineages (Marhold and Lihová 2006; Franzke, et al. 2011). Notably, the ancestors of the *Orychophragmus* genus underwent a unique WGD event (Zhang, et al. 2023), followed by rediploidization and speciation, resulting in a genome size of approximately 1.3 Gb—larger than that of other Brassicaceae species (Johnston, et al. 2005; Lysak, et al. 2008). Recently, *Orychophragmus* species have garnered attention as early-flowering ornamental plants and as promising industrial oilseed crops, owing to their high dihydroxy fatty acids content. This unique trait provides superior lubrication properties and ensures broad adaptability to diverse environmental conditions (Li, et al. 2018; Wang, et al. 2021; Huang, et al. 2023; Jia, et al. 2024). Moreover, their close evolutionary relationship with *Brassica* species—renowned for their ease of hybridization (Li and Heneen 1999; Li and Ge 2007)—makes *Orychophragmus* species valuable germplasm resources for *Brassica* genetics and breeding.

Despite their potential importance, the phylogenetic relationships within the *Orychophragmus* genus remain poorly understood. Currently, six species are recognized in this genus: *O. violaceus*, *O. longisiliqus*, *O. zhongtiaoshanus*, *O. taibaiensis*, *O. hupehensis*, and *O. diffusus* (Hu, et al. 2015; Hu, et al. 2016; Hu, et al. 2018). These species display significant morphological variation and inhabit diverse environments across East Asia (Fig. 1). However, previous analyses have relied on a limited number of nuclear and chloroplast genes, resulting in ongoing debates about the genus’s evolutionary history and relationships. To address these uncertainties, it is essential to incorporate broader genomic datasets and advanced methodologies to construct a more robust phylogeny, with particular attention to the potential effects of incomplete lineage sorting and introgression.

**Fig. 1.**
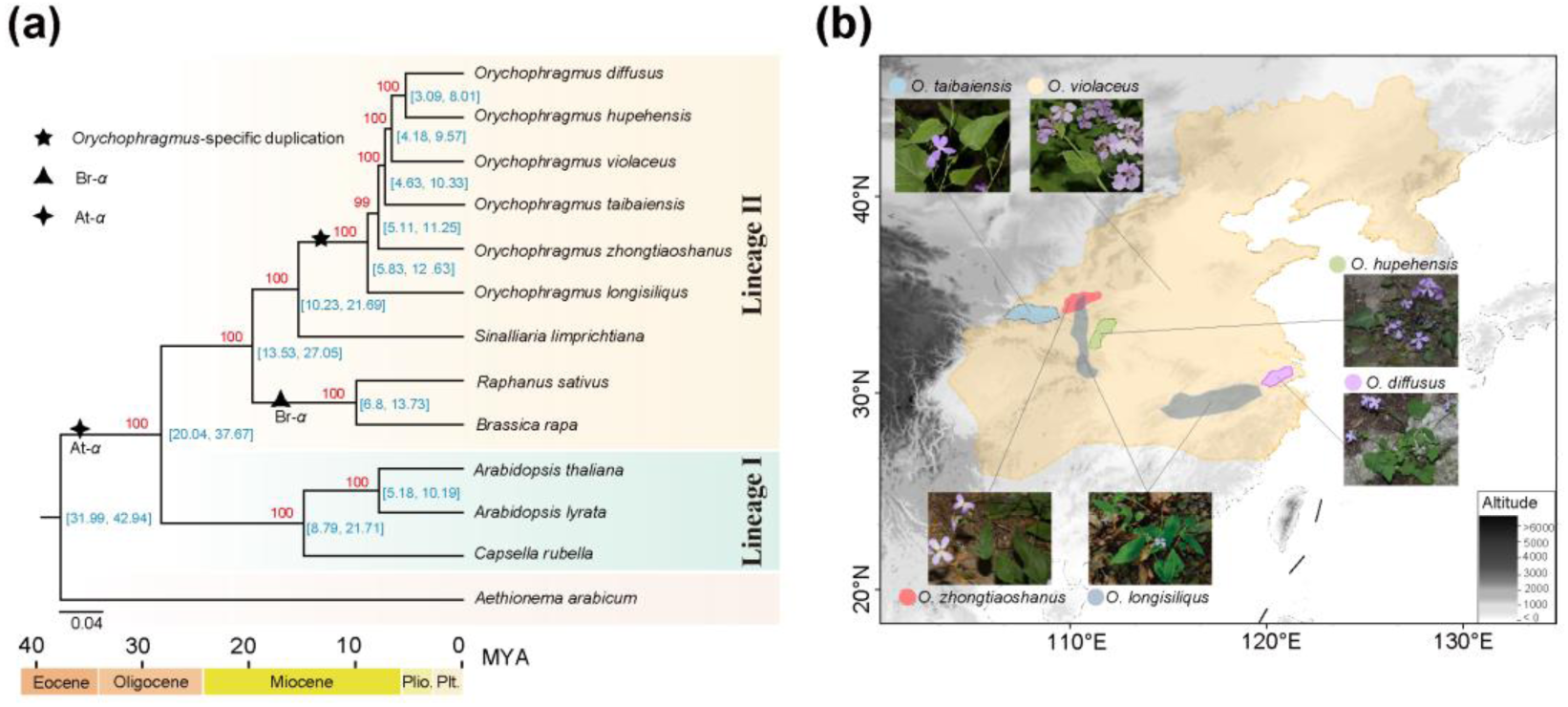
Phylogenetic relationships and geographical distribution of *Orychophragmus*. **(a)** Phylogenetic inference and divergence time estimation for six *Orychophragmus* species alongside other cruciferous species. Blue numbers indicate estimated divergence times (95% highest posterior density, in million years ago [MYA]), while red numbers represent bootstrap support values. The black star denotes the *Orychophragmus*-specific whole-genome duplication event, the black triangle marks the Brassica-specific triplication event, and the black tetragon represents the ancestral whole-genome duplication within the Brassicaceae family. **(b)** Geographical distribution and morphological characteristics of the six species within the genus *Orychophragmus*.

Notably, the *Orychophragmus* genus includes both widespread and regionally distributed species. *O. violaceus*, the most widely distributed species in the genus, is commonly cultivated as an ornamental plant and is known for its small purple flowers that typically bloom in early spring. In contrast, *O. taibaiensis* is endemic to the mountainous regions of the Taibai Mountains in northwest China. This endemic species is an ideal candidate for exploring its evolutionary and demographic histories, particularly its remarkable adaptation to the local alpine environment, in contrast to the more widespread and closely related *O. violaceus*. Additionally, *O. taibaiensis* has been reported to consist of both diploid (2n = 2x = 24) and tetraploid (2n = 4x = 48) plants (Zhou, et al. 2009), although the mode of origin of the polyploids remains unclear. As such, it also serves as an excellent model for studying the immediate evolutionary and genomic consequences of polyploidization. A direct comparison between tetraploid populations and their diploid progenitors could provide valuable insights into how polyploidization influences both the short-term adaptive responses and long-term evolutionary potential of these populations (Comai 2005; Mayrose, et al. 2011).

In this study, we integrated whole-genome sequencing and transcriptomic datasets to explore the phylogenetic relationships and divergence order of the six species within the genus *Orychophragmus*. We then examined the demographic and divergence history of the widespread species *O. violaceus* and the alpine endemic *O. taibaiensis*, comparing selection efficacy and the deleterious mutation load between the two species. Given that *O. taibaiensis* includes both diploid and tetraploid forms, we characterized the geographical distribution patterns of these cytotypes through extensive field investigations and karyotype analyses, while also investigating the origin of the tetraploid forms. Finally, we explored the genetic and gene expression changes associated with polyploidization in *O. taibaiensis* populations to gain deeper insights into the evolutionary consequences of whole-genome duplication and its potential short- and long-term selective effects on the ecology and evolution of these populations.

## 2 Materials and Methods

### 2.1 Taxon sampling, genomic sequencing and genetic data collection

After integrating newly sequenced genomic datasets from this study with transcriptomic datasets from previous research (Zhong, et al. 2019), we collected genomic and/or transcriptomic data for 52 individuals, representing all six *Orychophragmus* taxa: 7 *O. longisiliqus*, 13 *O. zhongtiaoshanus*, 10 *O. taibaiensis*, 1 *O. hupehensis*, 6 *O. diffusus*, and 14 *O. violaceus*. Additionally, we included one species of *Sinalliaria limprichtiana* as the outgroup. Detailed information about the data is provided in the supplemental information (Table S1). For the newly sequenced dataset, genomic DNA was extracted from leaf samples with the Qiagen DNeasy plant kit. Whole-genome paired-ends reads were generated using the T7 platform at BGI (PE150).

### 2.2 Phylogenomic construction

To construct the phylogenetic relationships among six *Orychophragmus* species and its relative species, we first downloaded protein sequences from six other species within the Brassicaceae family: *Aethionema arabicum*, *Capsella rubella*, *Arabidopsis lyrata*, *Arabidopsis thaliana*, *Raphanus sativus* and *Brassica rapa*. We then performed de novo transcriptome assembly using Trinity v.2.8.5 (Grabherr, et al. 2011) with default parameters applied to the filtered RNA-Seq reads processed by fastp v.0.20.1 (Chen, et al. 2018) for an individual from each species. We retained only the longest isoform for each gene and discarded redundant contigs using CD-HIT v4.8.1 (Fu, et al. 2012). Then, we utilized TransDecoder v.5.26.3 (Haas, et al. 2013) to identify protein-coding regions and subsequently used OrthoFinder v.2.5.4 (Emms and Kelly 2019) to recover single-copy genes. These single-copy genes were then used as references to extract the corresponding reads from each genome-resequenced individual and assemble each gene for each individual using aTRAM v.2.4.0 (Allen, et al. 2017). Finally, we used 514 single-copy genes longer than 300 bp for phylogenetic analysis. We aligned the amino acid sequences using MAFFT v7.475 (Katoh and Standley 2013) and then converted these alignments to codon alignments using PAL2NAL (Suyama, et al. 2006). After removing spurious sequences and poorly aligned regions with trimAl v1.4.rev22 (Capella-Gutiérrez, et al. 2009), the remaining alignments were combined into a supergene matrix for phylogenetic construction using RAxML v8.2.8(Stamatakis 2014) under the GTRCAT model. To estimate divergence times between species or clades, we employed the MCMCTree program implemented in the PAML package v4.10.0 (Yang 2007; dos Reis, et al. 2012), utilizing fossil data sourced from TimeTree (Kumar, et al. 2017). Furthermore, we performed coalescent-based species tree inference using ASTRAL v.5.15.5 (Yin, et al. 2019) and visualized gene trees and species tree discordances using DensiTree.v2.2.7 (Bouckaert 2010).

We further used population genomic data to cofirm the interspecific relationship within *Orychophragmus.* Whole-genome resequencing data and transcriptome data were combined to extract variant sites. For raw genomic resequencing and transcriptome sequencing reads, we used fastp v.0.20.1 (Chen, et al. 2018) for adapter removal and to trim bases from the start or end of reads if the base quality was <20. Reads shorter than 36 bases after trimming were further discarded. After quality control, all high-quality genomic resequencing reads were mapped to our new de novo assembled *O. violaceus* (Jia, et al. 2023) genome using the BWA-MEM algorithm of bwa v.0.7.17 (Li 2013) with default parameters, whereas the filtered transcriptome sequencing high-quality reads were mapped using HISAT2 v. 2.2.1 (Kim, et al. 2019). All generated bam files were subsequently sorted with SAMtools v.1.9 (Li, et al. 2009), and PCR duplicates were marked using the MarkDuplicates tool from Picard v.2.18.21 (http://broadinstitute.github.io/picard/, last accessed March 16, 2022). Genetic variants (SNP calling) were identified using the Genome Analysis Toolkit (GATK) v.4.2.5.0 (DePristo, et al. 2011) through its subcomponents HaplotypeCaller, CombineGVCFs, and GenotypeGVCFs. A merged VCF file including “all sites” (both variant and non-variant sites) were generated using the ‘EMIT_ALL_SITES’ flag. After combining the GVCF files from the resequencing and transcriptome datasets, the GenotypeGVCFs tool was applied to genotype the variants, which was followed by applying a strict set of filtering criteria: 1) Heng Li’s SNPable tool (http://lh3lh3.users.sourceforge.net/snpable.shtml) was used to mask genomic regions where reads were not uniquely mapped. To achieve this, the reference genome was divided into overlapping 100-mers, which were then aligned back to the genome using “bwa aln -R 1,000,000 -O 3 -E 3”. Only the sites (712,157,434 out of 1,285,736,775 sites) in which all 100-mers mapped uniquely and without mismatches were retained for downstream analyses; 2) SNPs with more than two alleles (>2), read depth (DP) < 5, or read depth greater than three times the average sequencing depth were removed; 3) SNPs with quality by depth (QD) < 2.0, Fisher Strand value (FS) >60, mapping quality (MQ) < 20, MQRankSum < -12.5 and ReadPosRankSum < -8.0 were filtered; 4) SNPs with a missing rate >20% were excluded. Finally, we used the Degeneracy tool to identify four-fold and zero-fold degenerate sites (https://github.com/tvkent/Degeneracy). Only four-fold degenerate sites were retained to create a robust intersect dataset after merging resequencing and transcriptome datasets with publicly available transcriptomic data from *Sinalliaria limprichtiana* (SRA ID: SRR6441722), which was used as an outgroup, and a total of 93,969 SNPs were used for population genomic phylogenetic constructions.

Neighbor-joining (NJ) phylogenetic trees were first constructed using PLINK v.1.90 (Purcell, et al. 2007) with the parameter distance 1-ibs to calculate the pairwise identify-by-state (IBS) genetic distance matrix, followed by MEGAZ (Kumar, et al. 2018) for tree construction. And maximum likelihood (ML) phylogenetic trees were constructed using IQ-TREE v.2.2.0.3 (Nguyen, et al. 2015). For the ML tree, ModelFinder (Kalyaanamoorthy, et al. 2017) was used to select the best-fitting substitution model (-B 1000 -m MFP) with 1000 ultrafast bootstraps. All trees were visualized using FigTree v1.4.4 (https://tree.bio.ed.ac.uk/software/Fig.tree). Additionally, to evaluate and test for the possibility of introgression among these *Orychophragmus* species, the ABBA-BABA tests were constructed using the Dtrios program in Dsuite (Malinsky, et al. 2021) to calculate the Patterson’s *D*-statistics and corresponding test values (*Z-*scores) (Patterson, et al. 2012). The *D*-statistics (*D*) were useful to detect introgression, while the *Z*-scores (*Z*) were used to assess statistical significance, with an absolute value of Z > 3 often considered as a critical threshold for gene flow (Excoffier, et al. 2013; Martin, et al. 2015).

### 2.3 Demographic history analyses of *O. violaceus* and *O. taibaiensis*

To reconstruct the historical demography of the widespread species *O. violaceus* and the locally endemic species *O. taibaiensis*, species distribution models (SDMs) (Elith and Leathwick 2009) were developed using MAXENT v.3.4.4 (Phillips and Dudík 2008; Merow, et al. 2013). These models were applied to evaluate shifts in suitable habitat ranges across different time periods for both species. To achieve this, current native distribution records for *O. violaceus* (445 records) and *O. taibaiensis* (91 records) were collected from the Chinese Virtual Herbarium (CVH, https://www.cvh.ac.cn/) and field investigations. Environmental layers for 19 biologically significant climate variables (BIO1–BIO19) at a 2.5-arc-minute resolution were obtained from the WorldClim database (http://www.worldclim.org) (Hijmans, et al. 2005) for three time periods: the present (1970–2000), the Mid Holocene (∼6,000 years ago), and the Last Glacial Maximum (∼20,000 years ago). To minimize overfitting, only climate variables with Pearson correlation coefficients below 0.8 (r < 0.8) were included in the SDM analysis. The accuracy of the SDMs was evaluated using the area under the curve (AUC) of the receiver operating characteristic (ROC) plot generated by Maxent (Fawcett 2006), which is a threshold-independent and prevalence-insensitive metric.

To further investigate the demographic history of the two species, we initially applied the Pairwise Sequentially Markovian Coalescent (PSMC) method (Li and Durbin 2011) to infer the historical effective population size (*N_e_*) dynamics. The analysis was conducted using the parameters ‘N25 -t15 -r5 -p “4+25×2+4+6”’ for both species, with 100 bootstrap replicates performed. Assuming a mutation rate of 8.22 × 10^-9^ mutations per site per generation (Beilstein, et al. 2010) and a generation time of 1 year, we converted the coalescent-scaled time into absolute time measured in years. To gain deeper insights into recent population dynamics, particularly over the past 10,000 years, we employed the Multiple Sequentially Markovian Coalescent approach (MSMC2) (Schiffels and Durbin 2014). This analysis was performed on phased whole-genome sequences that were generated by Beagle v.4.1 (Browning and Browning 2009) from four individuals (representing eight haplotypes) for each species. For both species, MSMC2 was run on all possible individual configurations, and the medians and standard deviations of *N_e_* changes were subsequently estimated.

Subsequently, we inferred the divergence history between *O. violaceus* and *O. taibaiensis* using a coalescent simulation-based approach implemented in *fastsimcoal*2 v.27 (Excoffier, et al. 2013; Excoffier, et al. 2021). From the whole-genome resequencing dataset of the two species, we utilized only fourfold degenerate sites to construct the two-dimensional site frequency spectrum (SFS) via easySFS (https://github.com/isaacovercast/easySFS). We evaluated twelve demographic models, each representing different scenarios, including divergence with or without gene flow, and constant or variable population sizes following divergence (Fig. S6). For each model, we performed 50 independent runs, with each run consisting of 40 cycles and 100,000 coalescent simulations. Model comparisons were based on the maximum likelihood values from the 50 runs, further assessed using Akaike weights and Δlikelihood. The model with the highest Akaike weight was selected as the optimal demographic scenario. To estimate confidence intervals, we conducted parametric bootstrapping with 100 bootstrap replicates, each including 50 independent runs. As in the PSMC and MSMC2 analyses, we assumed a mutation rate of 8.22 × 10^-9^ mutations per site per generation (Beilstein, et al. 2010) and a generation time of 1 year.

### 2.4 Assessment of selection efficiency and divergent selection signatures between O. violaceus and O. taibaiensis

Given the differences in demographic history and effective population sizes between *O. violaceus* and *O. taibaiensis*, we compared the efficacy of natural selection and the deleterious genetic load between the two species. First, we calculated and compared the genome-wide ratio of 0-fold nonsynonymous to 4-fold synonymous nucleotide diversity (π_0-fold_/π_4-fold_) using pixy v1.0.4 (Korunes and Samuk 2021) across 100-kb nonoverlapping window. Next, we used the folded site frequency spectrum (SFS) of 0-fold nonsynonymous and 4-fold synonymous sites to estimate the distribution of fitness effects (DFE) for new nonsynonymous mutations using DFE-alpha v2.16 (Keightley and Eyre-Walker 2007). A demographic model with a stepwise population size change was fitted to the neutral SFS, and the estimated parameters were incorporated to infer the fitness effects of new deleterious mutations and the strength of purifying selection (*N_e_s*) for each species. Confidence intervals (95% CIs) for all estimates were generated using 200 bootstrap replicates, resampling across all sites within each class and excluding the top and bottom 2.5% of replicates.

Furthermore, to investigate the biological significance of genomic regions potentially under divergent selection during the divergence of *O. violaceus* and *O. taibaiensis*, we identified candidate divergent selection regions using the XP-CLR method (Chen, et al. 2010). Specifically, we calculated cross-population composite likelihood ratio (XP-CLR) scores between the two species within 5-kb non-overlapping windows. Genomic regions with scores in the top 1% (2397 windows) were identified as candidate selective regions. To validate these regions, we calculated independent estimates of inter-species genetic divergence (*F*_ST_) and intra-species Tajima’s *D* values, confirming signatures of divergent selection. Finally, we extracted the genes (340) located within the candidate regions and performed Gene Ontology (GO) enrichment analysis using the topGO R package (https://bioconductor.org/packages/topGO/) to investigate the biological significance of these regions.

### 2.5 Determination of diploid and tetraploid distribution across *O. taibaiensis* populations

To thoroughly investigate the geographical distribution of diploid and tetraploid cytotypes across the natural populations of *O. taibaiensis*, seeds were collected from 94 individuals across seven populations, covering the entire known distribution of *O. taibaiensis* identified in the field. The seeds were germinated on wet filter paper in Petri dishes at 25°C. Root meristems were carefully excised from germinating seeds and pre-treated with a 0.1% colchicine solution at 25°C for 3 hours. The samples were then washed and fixed in a 3:1 solution of ethyl alcohol and acetic acid at 4°C for 3 hours. Subsequently, the samples were rinsed in distilled water and immersed in 1 M HCl at 37°C for 45 minutes to facilitate hydrolysis. After hydrolysis, the root tips were washed in distilled water and stained with a modified carbol-fuchsin solution at room temperature for more than 3 hours. Finally, the meristems were dissected and squashed onto glass slides, and chromosomal counts were analyzed using a microscope under oil immersion (×1000 magnification) and photographed.

To further investigate and differentiate the origin of polyploidy (autopolyploidy or allopolyploidy) in the identified diploids and tetraploids of *O. taibaiensis*, we selected five diploids and five tetraploids for high-depth whole-genome resequencing (average depth ∼52.48×). First, we utilized nQuire (Weiß, et al. 2018) to confirm the ploidy levels of the sequenced individuals. The “denoise” subcommand was applied to reduce baseline noise, and sequencing reads were mapped back to genome assemblies to evaluate read depths and allele frequencies. The density distribution of the three allele types (reference, alternative, and both) was calculated to validate ploidy levels. Next, we integrated GenomeScope and Smudgeplots (Ranallo-Benavidez, et al. 2020) to distinguish between autotetraploids and allotetraploids. This was achieved by analyzing patterns of nucleotide heterozygosity for polyploid samples, following the construction of *K-mer* frequency distributions using Jellyfish v.2.2.9 (Marçais and Kingsford 2011) from the resequencing data. Finally, we re-performed variant calling for the five resequenced individuals of tetraploid *O. taibaiensis* using GATK, setting the parameter “--ploidy 4”. The relative proportions of different genotypes (e.g., Aaaa, AAaa, aaaA) were calculated to further verify whether the tetraploids were of autotetraploid or allotetraploid origin.

### 2.6 Population genomic analysis and assessment of genetic load for diploid and tetraploid individuals of *O. taibaiensis*

To construct the phylogenetic relationships of *O. taibaiensis* at different ploidy levels, we re-performed variant calling and filtering, following the method described in section 2.2, for the ten high-depth resequenced individuals. First, we used PLINK v.1.90 (Purcell, et al. 2007) with the parameter distance 1-ibs to calculate the pairwise identify-by-state (IBS) genetic distance matrix, quantifying the relatedness among the five diploids and five tetraploids. Based on the resulting distance matrix, we employed MEGAZ (Kumar, et al. 2018) to construct an unrooted NJ tree among individuals with different ploidy levels. Next, we utilized *fastsimcoal2* (Excoffier, et al. 2021) to infer the divergence history of the two cytotypes based on a two-dimensional joint SFS of fourfold degenerate sites. Whole-genome resequencing data from *O. longisiliqus* and *O. zhongtiaoshanus* were included as outgroups to infer the derived states of alleles using the est-sfs software (Keightley and Jackson 2018). The analysis was performed under two models: one accounting for gene flow and one assuming no gene flow (see Fig. S14a, S14b). The detailed procedures followed those outlined in section 2.3.

To test for the potential effects of polyploidization on the efficacy of purifying selection and the accumulation of deleterious genetic load, we estimated and compared the ratio of 0-fold nonsynonymous to 4-fold synonymous nucleotide diversity (π_0-fold_/π_4-fold_) between diploid and tetraploid individuals of *O. taibaiensis*. Additionally, we examined and compared the distribution of fitness effects (DFE) of new mutations across both ploidy levels. The intensity of purifying selection (*N_e_s*) at 0-fold nonsynonymous sites was estimated using DFE-alpha v2.16 (Keightley and Eyre-Walker 2007) by fitting a stepwise population size change model to account for differences in the SFS of 0-fold nonsynonymous sites and putatively neutral 4-fold synonymous sites. To evaluate the potential effects of the diploid assumption in these analyses, we re-performed variant calling for tetraploid individuals using GATK, setting the parameter "--ploidy 4". For each tetraploid individual, we randomly subsampled two alleles from the four alleles per site to generate six independent subsampling datasets. These datasets were then used for associated analyses, including *fastsimcoal2* modeling, genetic load estimation, and DFE analysis. This approach ensured robust comparison between diploid and tetraploid cytotypes while accounting for the polyploid nature of the tetraploid individuals.

### 2.7 Differential gene expression analysis between the two cytotypes of *O. taibaiensis*

To investigate how polyploidy influences transcriptomic changes, seeds of diploid and tetraploid *O. taibaiensis* collected from two geographic regions (Fig. 3a red triangles) were grown under identical conditions in a growth chamber. The environment was controlled with a 16/8 h light/dark photoperiod, a constant temperature of 25°C, and a light intensity of 100 μmol m⁻² s⁻¹. After six months of growth, leaf and root tissues were collected from plants with similar growth characteristics for both cytotypes. Three biological replicates were collected per cytotype. Harvested materials were frozen in liquid nitrogen and stored at -80°C until RNA extraction. Total RNA was extracted using the TIANGEN RNA Kit following the manufacturer’s protocol. Sequencing libraries were constructed and sequenced on the DNBSEQ-T7 platform using paired-end sequencing strategy.

After filtering and trimming RNA reads using fastp v.0.20.1 (Chen, et al. 2018) with default parameters, the clean reads were mapped to the *O. violaceus* genome (Jia, et al. 2023) using HISAT2 v.2.2.1 (Kim, et al. 2019). Gene expression levels were quantified with StringTie v1.3.6 (Pertea, et al. 2015; Kovaka, et al. 2019), and normalized expression levels were calculated as transcripts per million (TPM). Out of 52,812 annotated genes in the reference genome (Jia, et al. 2023), a total of 37,607 genes (TPM > 0 for each sample) were expressed across both leaf and root tissues in diploid and tetraploid samples. For differential expression analysis, the R package *DESeq2* (Love, et al. 2014) was used, with thresholds of |log2 fold change (FC)| ≥ 2 and adjusted *p*-value < 0.001 to identify significant differentially expressed genes (DEGs). Genes with |log2 fold change (FC)| < 1 or adjusted *p*-value > 0.05 were classified as non-differentially expressed genes (NDEs). The DEGs were further clustered and visualized using the ClusterGVis package (https://github.com/junjunlab/ClusterGVis), which employs the “mfuzz” clustering method to group genes based on their expression patterns. To identify the biological functions enriched among the DEGs associated with polyploidy in *O. taibaiensis*, GO enrichment analysis was performed using the topGO R package (https://bioconductor.org/packages/topGO/).

### 2.8 Selection on genes exhibiting differential expression between cytotypes

To examine and compare differences in the strength and direction of natural selection on DEGs and NDEs associated with polyploidization in diploids and tetraploids of *O. taibaiensis*, we first estimated pairwise relative genetic divergence (*F*_ST_), and absolute genetic divergence (*d_xy_*) within and between resequenced individuals of different cytotypes for the DEGs and NDEs gene sets using PIXY v1.0.4 (Korunes and Samuk 2021). Next, we calculated the ratio of zero-fold nonsynonymous to four-fold synonymous nucleotide diversity to compare deleterious mutation loads between the two gene sets. Finally, we assessed the strength of selective constraint on the two gene sets by modeling the distribution of fitness effects (DFE). As described earlier, the strength of purifying selection (*N*_e_s) on zero-fold nonsynonymous sites was estimated for each gene set in both diploids and tetraploids, using four-fold synonymous sites as a neutral reference, with DFE-alpha v2.16 (Keightley and Eyre-Walker 2007). For tetraploid individuals, six additional independent datasets, generated by randomly sampling two alleles from the four alleles, were applied in all analyses in this section.

## 3 Results

### 3.1 Phylogenetic and evolutionary relationships among species in the genus *Orychophragmus*

To investigate the phylogenetic relationships within the genus *Orychophragmus*, we incorporated seven additional species from the Brassicaceae family and utilized 514 single-copy genes to construct phylogenetic trees using both concatenated and coalescent approaches. Phylogenetic analysis based on single-copy genes revealed the monophyly of *Orychophragmus*, with its closest relative being the genus *Sinalliaria* (Fig. 1a). According to MCMCTree, the divergence between *Orychophragmus* and its closest relative *Sinalliaria* was estimated to be 15.43 million years ago (MYA, 95% highest posterior density: 10.23-21.69 MYA). Within the clade *Orychophragmus*, the concatenated analysis produced a strongly supported topology of (outgroups, (*O. longisiliqus*, (*O. zhongtiaoshanus*, (*O. taibaiensis*, (*O. violaceus*, (*O. diffusus*, *O. hupehensis*)))))) (Fig. 1a). This topology was also supported by the ASTRAL coalescent approach (Fig. S1a), although some branches exhibited weak support values. To further explore this, we used DensiTree as a visualization tool to display and quantify the concordance and discordance between individual gene trees and the species tree, which revealed significant inconsistencies between many individual gene trees and the species tree within the *Orychophragmus* clade (Fig. S1b).

To further validate the phylogenetic relationship within the genus *Orychophragmus*, we integrated population-level resequencing and transcriptomic datasets, using *Sinalliaria* as the outgroup to construct phylogenies. The phylogenetic relationships within *Orychophragmus*, inferred using both maximum likelihood (ML) and neighbor-joining (NJ) methods (Fig. S2), were consistent with those derived from single-copy gene analyses (Fig. 1a, Fig. S1). Given the observed inconsistencies in individual gene trees within *Orychophragmus*, we used ABBA-BABA analyses to assess potential gene flow between these species. Specifically, we systematically tested for signatures of introgression by calculating Patterson’s *D*-statistics and corresponding test values (*Z*-scores) (Patterson, et al. 2012). The results provide strong evidence of pervasive historical introgression among the *Orychophragmus* species (Fig. S3), which could partially explain the observed inconsistencies in the individual gene trees of *Orychophragmus*.

### 3.2 Demographic histories and divergent selection between *O. violaceus* and *O. taibaiensis*

Given the unique features of *O. taibaiensis*—the only species in the genus reported to consist of both diploids and tetraploids as well as being the endemic species found at the highest altitudes within the genus (Fig. S4)—we specifically explored and compared the demographic histories of this species with the closely related widespread species *O. violaceus*. To achieve this, we first applied species distribution models (SDMs) to reconstruct the suitable habitats of *O. violaceus* and *O. taibaiensis* across three evolutionary periods: the present, the Middle Holocene (MH, ∼6 ka), and the Last Glacial Maximum (LGM, ∼21–18 ka). Our analysis revealed that the suitable habitat of *O. violaceus* expanded significantly from the LGM and MH to the present. In contrast, the suitable habitat of *O. taibaiensis* remained relatively stable, being primarily confined to the Qinling-Daba mountain regions (Fig. 2a). Furthermore, over time, its range became increasingly restricted to the Taibai Mountain region, from the LGM and MH to the present.

**Fig. 2.**
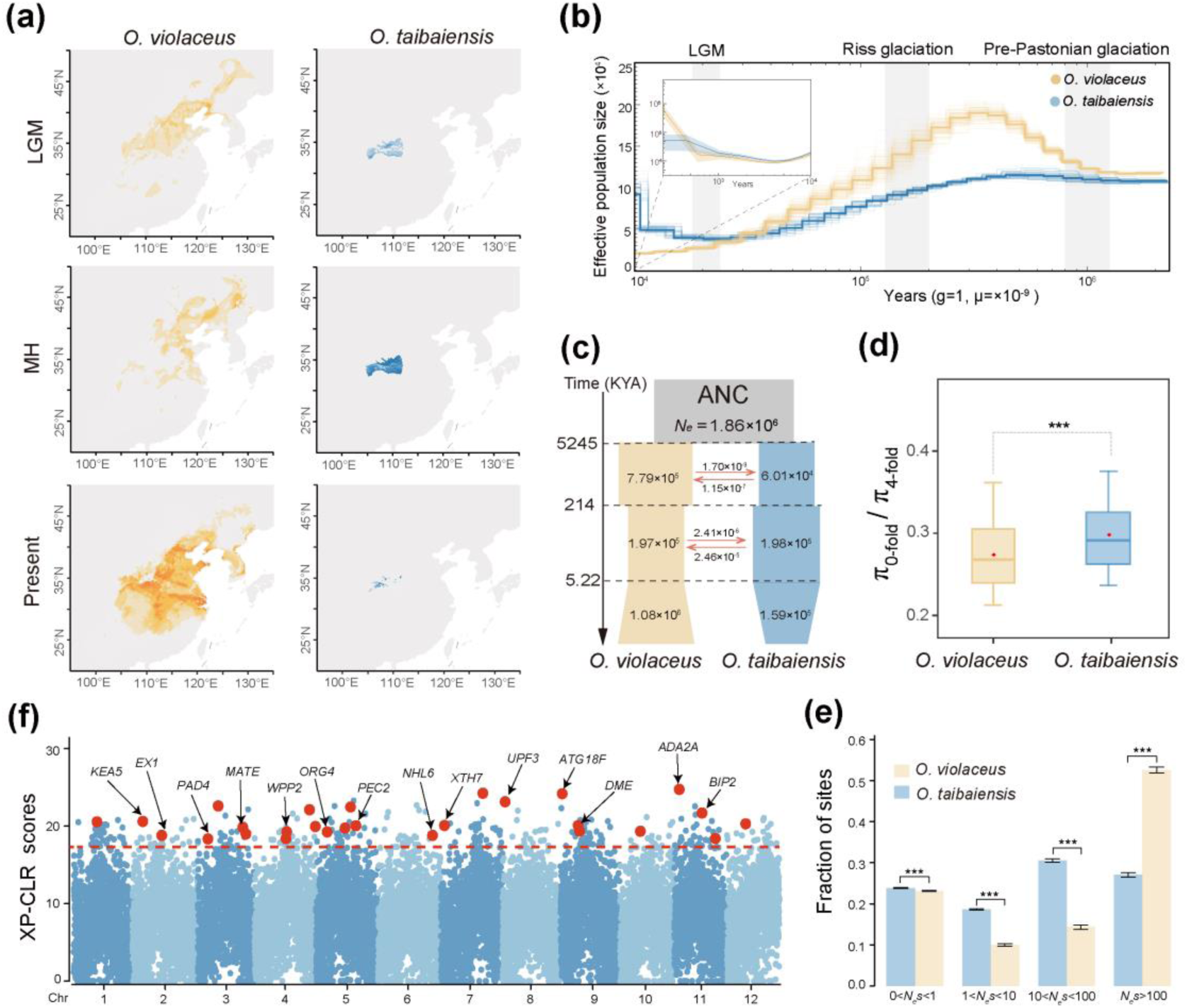
Demographic histories and divergent selection between *O. violaceus* and *O. taibaiensis*. **(a)** Species distribution modeling under the Last Glacial Maximum (LGM), Mid-Holocene (MH), and present climate conditions. **(b)** Demographic history inferred using the PSMC (outer) and MSMC (inner, see Fig. S5 for details) model. Gray vertical bars indicate the LGM, Riss glaciation, and Pre-Pastonian glaciation periods. Bold lines represent dynamic changes in effective population size (*N*_e_), while faint lines show 100 bootstrap replicates, ensuring robustness. **(c)** The best-fit demographic model inferred using *fastsimcoal2*. Each block represents a current or ancestral population, with arrows indicating gene flow after divergence (per-generation migration rates). The timing of historical events is shown in thousands of years ago (KYA). **(d)** Ratio of nucleotide diversity at 0-fold sites relative to 4-fold sites. **(e)** Distribution of fitness effects (DFE) in bins of *N*_e_s for new 0-fold nonsynonymous mutations for *O. violaceus* and *O. taibaiensis*. Error bars indicate 95% confidence intervals based on 200 bootstrap replicates. **(f)** Selective sweep analysis based on XP-CLR scores along chromosomes. The top 1% of scores, above the red dashed horizontal line, are considered candidate selective regions. Red circles highlight representative candidate genes located within these regions, with black arrows indicating their names. Asterisks indicate statistical significance from the Wilcoxon test (two-tailed) (NS. > 0.05, \**P* < 0.05, \*\**P* < 0.01, \*\*\**P* < 0.001).

**Fig. 3.**
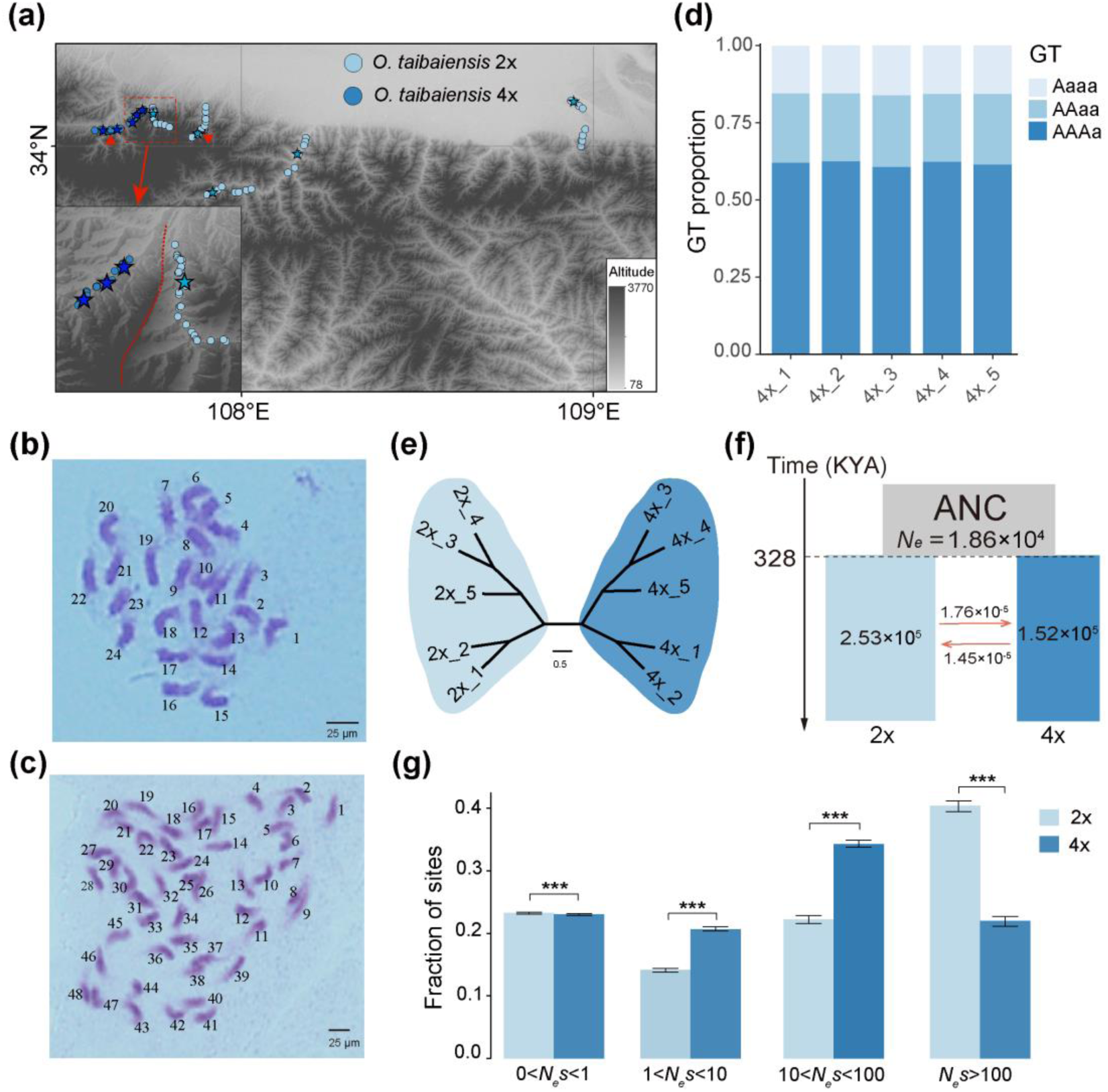
Comparison of the geographic distribution, divergence history, genetic load, and selection efficiency of diploid and tetraploid *O. taibaiensis*. (a) Geographic distribution of diploid and tetraploid *O. taibaiensis*, with stars representing the sequenced individuals mentioned in the text. Red triangles indicate the seed collection sites used for gene expression analysis across ploidy levels. (b) Karyotype of diploid *O. taibaiensis*. (c) Karyotype of tetraploid *O. taibaiensis*. (d) Relative proportions of various genotypes within the five whole-genome resequenced tetraploid *O. taibaiensis* individuals. (e) Phylogenetic relationships among the five resequenced diploid and tetraploid *O. taibaiensis*. (f) Population historical dynamics and divergence time estimation between diploid and tetraploid *O. taibaiensi*s, based on *Fastsimcoal2*. (g) Distribution of fitness effects (DFE) in bins of *N*_e_s for new 0-fold nonsynonymous mutations for diploid and tetraploid *O. taibaiensis*. Error bars represent 95% confidence intervals based on 200 bootstrap replicates. Asterisks indicate the level of significance in the Wilcoxon test (two-tailed) (NS: not significant, \**P* < 0.05, \*\**P* < 0.01, \*\*\**P* < 0.001).

To assess the long-term effective population size (*N*_e_) dynamics of these two species, we applied the pairwise sequential Markovian coalescent (PSMC) method (Li and Durbin 2011). The results revealed that both species experienced a reduction in *N*_e_ from the Riss glaciation to the LGM (Fig. 2b). Consistent with the broader distribution range of *O. violaceus* predicted by the SDMs, the *N*_e_ of *O. violaceus* was generally much larger than that of *O. taibaiensis* (Fig. 2b). Since PSMC can only estimate population dynamics up to around 10,000 years ago, we also employed MSMC2 to focus specifically on the population histories of these two species over the latest 10,000 years. The results indicated that the *O. violaceus* population underwent a more significant recovery following LGM when compared to *O. taibaiensis* (Fig. 2b; Fig. S5).

To further infer the divergence history of the two species, we employed a coalescent simulation-based approach using *fastsimcoal*2 (Excoffier, et al. 2013; Excoffier, et al. 2021). Twelve models were evaluated (Fig. S6), differing in the presence or absence of post-divergence gene flow and changes in population size following species divergence. The best-fitting model (Model 12 in Fig. S6; Table S2) suggested that *O. violaceus* and *O. taibaiensis* diverged approximately 5.25 MYA (95% confidence interval (CI) = 1.40–7.33 MYA) (Fig. 2c; Table S3), consistent with the divergence time estimated from phylogenetic analysis (Fig. 1a). Furthermore, the model revealed divergent patterns of changes in effective population sizes and asymmetric gene flow between the species following their divergence, although no gene flow has occurred between the two species within the recent 5.22 thousand years ago (KYA) (95% CI = 5.02–5.79KYA) (Fig. 2c, Table S2). Additionally, the model inferred that *O. taibaiensis* experienced a recent population reduction (*N_e_* = 1.59×10^5^, [1.45×10^5^–1.99×10^5^]), in contrast to *O. violaceus*, which underwent population expansion in both effective population size (*N_e_* = 1.08×10^6^ [8.45×10^5^–1.29×10^6^]) and distribution range (Fig. 2a,c).

To further compare the likely influence of different demographic histories on the efficacy of selection and the accumulation of mutational load between the two species, we estimated the ratio of nucleotide diversity at 0- to 4-fold degenerate sites. Consistent with the expectation that species with smaller population sizes have a reduced efficacy of selection to purge deleterious mutations at 0-fold sites (Lynch, et al. 1995), we observed that the ratio of 0- to 4-fold nucleotide diversity was significantly elevated in *O. taibaiensis* compared to *O. violaceus* (Fig. 2d). Similarly, the distribution of fitness effects (DFE) analysis revealed that *O. taibaiensis* exhibited weaker purifying selection against strongly deleterious nonsynonymous mutations (*N_e_s* > 100) compared to *O. violaceus* (Fig. 2e). Moreover, given that *O. taibaiensis* contains both diploid and tetraploid cytotypes, we re-analyzed and compared the population history divergence, genetic load, and DFE solely between *O. violaceus* and diploid *O. taibaiensis*. These results were highly consistent with the broader analysis, further supporting the conclusion that *O. taibaiensis* exhibits reduced selection efficacy and a higher mutational load compared to *O. violaceus* (Fig. S7b).

Lastly, we applied and calculated XP-CLR statistics to identify potential divergent selection regions between the two species. In total, we detected 2,397 outlier windows in the top 1% of XP-CLR scores (Fig. 2f). Further gene ontology (GO) enrichment analyses of genes within the candidate selective regions revealed significant enrichment in GO terms such as “cellular macromolecule metabolic process” and “response to reactive oxygen species” (Fig. S8a; Table S4), suggesting their association with divergent adaptation to different altitudinal environments between the two species. Many genes known to be involved in stress and defense responses were identified within these regions. For example, the *Arabidopsis* orthologous gene *PAD4*, which plays a critical role in salicylic acid signaling and resistance gene-mediated plant disease resistance(Zeng, et al. 2023; Yu, et al. 2024), was found within these regions. In addition to the significant signals of XP-CLR, we observed substantially increased inter-species genetic divergence (*F*_ST_) and reduced intra-species Tajima’s *D* values (Fig. S8b), providing strong evidence of divergent selection within these genic regions. Similarly, the *Arabidopsis* orthologous gene *PEC2*, which is involved in responding to environmental stimuli such as jasmonic acid, light, and wounding (Völkner, et al. 2021), was also identified, highlighting its role in stress responses and adaptive signaling pathways. We found substantially increased *F*_ST_ and reduced Tajima’s *D* values in both species around these genic regions (Fig. S8b). Similar patterns were observed for other genes related to fundamental processes crucial to plant growth, development, and epigenetic regulation, such as genes *DME* (Khouider, et al. 2021) and *XTH7* (Volyanskaya, et al. 2023) (Fig. S8d,e)

### 3.3 Cytotype diversity and distribution in natural populations of *O. taibaiensis*

Although *O. taibaiensis* was previously reported to have both diploid and tetraploid cytotypes locally distributed in the Taibai Mountains of central China (Tan, et al. 1998; Zhou, et al. 2008), the spatial overlap and distribution patterns of populations with different ploidy levels remain unclear. To address this, we determined the ploidy levels of *O. taibaiensis* in a relatively large sample set of 94 individuals from various locations. Chromosome number determination confirmed the findings of earlier studies (Zhou, et al. 2008) and revealed variation in chromosome size. Specifically, we identified two ploidy levels, with chromosome counts of 2n = 24 for diploids (Fig. 3b) and 2n = 48 for tetraploids (Fig. 3c). Cytotype distribution analysis indicated that individuals with different ploidy levels were primarily separated by local mountain barriers within relatively limited geographical ranges (Fig. 3a). To further investigate whether there was environmental differentiation among cytotypes, we performed a principal component analysis (PCA) using five uncorrelated environmental variables derived from 19 bioclimatic factors (bio1-bio19) and elevation (Fig. S9a,c, Table S5, Table S6). Our results revealed little to no evidence of niche differentiation between cytotypes, with only subtle variation observed along environmental and climatic gradients (Fig. S9).

Next, we selected five diploid and five tetraploid individuals of *O. taibaiensis* for high-depth whole-genome resequencing. We first employed nQuire (Weiß, et al. 2018), a statistical method designed to determine the most plausible ploidy model based on the distribution of base frequencies in the sequencing data. The results from nQuire confirmed the diploid and tetraploid forms of *O. taibaiensis* that we had previously identified (Fig. S10a, b). Specifically, the read depth density distribution of the three alleles (alternative, reference and both) in diploid individuals was close to 1:1:2 (Fig. S10c), while in tetraploid individuals, it was approximately 1:3:4 (Fig. S10d). To differentiate between autopolyploidization and allopolyploidization, we utilized Smudgeplots (Ranallo-Benavidez, et al. 2020) to visualize the expected allele ratio patterns (Fig. S11). We found that the AAAB pattern was more prominent than AABB for tetraploids, consistent with the expectation of an autopolyploid origin. Additionally, genome-wide heterozygosity analysis using GenomeScope v.2.0 revealed a distribution of aaab (3.52%) > aabb (1.96%) (Fig. S12a), further supporting the notion of autotetraploids. Lastly, we recalled SNPs based on the tetraploid model for each of the five resequenced tetraploids and calculated the proportions of various genotypes. The results indicated that the AAAa genotype was significantly more prevalent than AAaa across the entire genome (Fig. 3d; Fig. S12b-f), reinforcing the hypothesis of autotetraploidy in the polyploid *O. taibaiensis*.

### 3.4 Origin history of autotetraploids and the evolutionary consequences of short-term polyploidization in *O. taibaiensis*

To investigate the origin and genetic differentiation between diploid and autotetraploid *O. taibaiensis*, we constructed an unrooted tree using ten highly deep-sequenced individuals from both ploidy levels. The analysis revealed that the individuals formed two distinct clusters, albeit with short genetic distances and low divergence (Fig. 3e). Using *fastsimcoal2*, we identified the gene flow model as the best fit for the divergence history and estimated that diploids and tetraploids diverged approximately 328 KYA (95% CI: 326–468 KYA). The contemporary *N*_e_ of diploids was estimated to be 2.53×10^5^ (95% CI: 2.40×10^5^–3.61×10^5^), which is slightly larger than that of autotetraploids, estimated at 1.52×10^5^ (95% CI: 1.45×10^5^–2.06×10^5^) (Fig. 3f, Table S7).

To further explore the effects of ploidy on the efficacy of purifying selection, we evaluated and compared the DFE between diploids and tetraploids. The results showed that diploids exhibited a significantly higher proportion of loci under strong purifying selection (*N*_e_s > 100) compared to autotetraploids. This finding suggests relaxed purifying selection acting on nonsynonymous sites in the tetraploid population compared to the diploid population (Fig. 3g). These results remained robust when we randomly subsampled two out of four alleles per site from the genotypes of tetraploid population to account for potential biases introduced by genotype calling (Fig. S13). However, it is important to note that such biases cannot be fully resolved, as a diploid model was assumed for DFE estimation. Additionally, we calculated the ratio of overall 0- to 4-fold nucleotide diversity (Fig. S14c-S14i) and found that the ratio was significantly higher in autotetraploids compared to diploids. These results further suggest that the tetraploid population has accumulated a higher proportion of deleterious mutations, which aligns with the findings from *fastsimcoal2* and DFE analysis, showing that the tetraploid population, with higher genetic load, exhibits a smaller effective population size and lower purifying selection efficiency.

### 3.5 Gene expression changes and associated signatures of selection upon polyploidization in *O. taibaiensis*

To explore transcriptional changes and responses to WGD, we examined gene expression differences using RNA-seq data from diploids and autotetraploids of *O. taibaiensis* in both leaf and root tissues. Principal component analysis (PCA) clearly distinguished samples based on tissue type and cytotype (Fig. S15a). To specifically investigate gene expression changes associated with polyploidization, we identified differentially expressed genes (DEGs) between diploids and autotetraploids. While the majority of genes showed no significant change in expression (NDEs), we identified 3,254 and 3,422 DEGs in leaf and root tissues, respectively (Fig. S15b), with 1,910 DEGs shared between the two tissues.

GO enrichment analysis of these DEGs revealed that genes differentially expressed between diploids and tetraploids are highly enriched in terms related to signal transduction, cell communication, defense response, regulation of gene expression, and circadian rhythm (Fig. 4a). For instance, genes downregulated in tetraploids, such as *SNI1* and *CERK1*, are involved in gene transcription, DNA recombination, and defense signaling (Fig. 4b). Notably, *SNI1* in *Arabidopsis* plays a crucial role in preventing errors during meiotic recombination in plants (Liu, et al. 2018; Zhu, et al. 2021). Similarly, *α-DOX1* and *ADC* are key players in metabolic processes, with *α-DOX1* participating in fatty acid alpha-oxidation and *ADC* being involved in polyamine biosynthesis, both of which are essential for maintaining cellular integrity and stress responses (Vicente, et al. 2012; Rossi, et al. 2015) (Fig. 4b). The significantly reduced expression of these genes in tetraploids may be attributed to genomic instability and the regulatory complexity caused by gene dosage effects in tetraploids. Conversely, in autotetraploid plants, the upregulated genes are primarily involved in functions critical for maintaining genomic stability and responding to environmental stresses. For example, *EXT3* is essential for plant cell wall organization, while *RECA2* is involved in DNA repair (Cannon, et al. 2008; Saha, et al. 2013; Odahara, et al. 2017). Additionally, *LecRK* and *LNK3* have been reported to play roles in cellular responses to salicylic acid, defense against pathogens, and the regulation of circadian rhythms (Bouwmeester and Govers 2009; Kidokoro, et al. 2023). These findings may reflect a broader theme of adaptation and stress response, highlighting the ability of tetraploids to cope with environmental challenges (Fig. 4a, 4b).

**Fig. 4.**
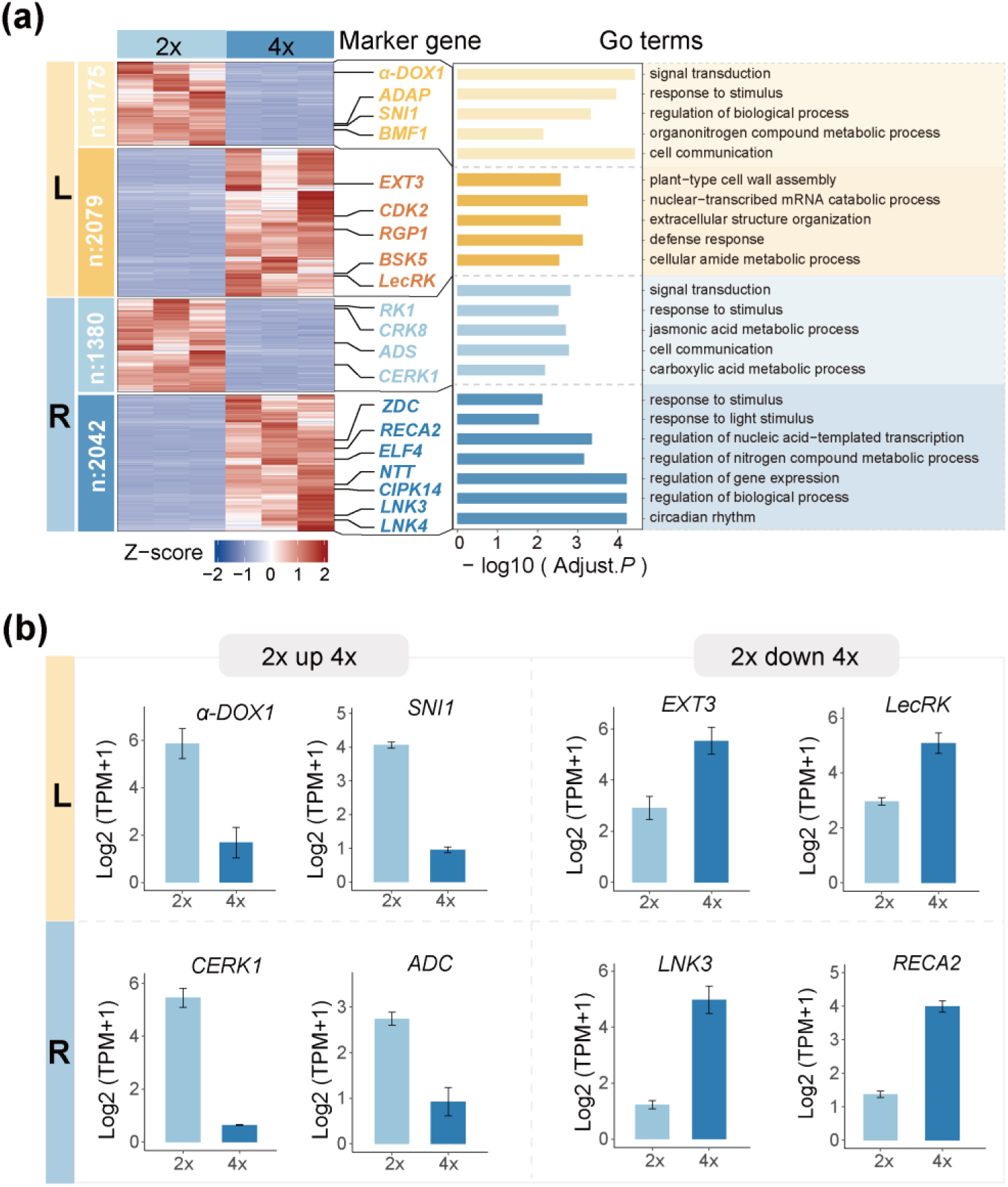
Differential expression analysis between diploid and tetraploid *O. taibaiensis*. (a) Hierarchical clustering of gene expression in diploid and autotetraploid *O. taibaiensis* for leaves (yellow, L) and roots (blue, R), highlighting key representative Gene Ontology (GO) terms enriched among differentially expressed genes (DEGs) within each cluster. Multiple representative DEGs from each cluster are shown. (b) Examples of expression level comparisons between diploid and tetraploid *O. taibaiensi*s for selected representative DEGs identified in (a).

To further investigate the association between expression divergence and sequence divergence following genome doubling in *O. taibaiensis*, we compared both the relative (*F_ST_*) and absolute (*d_xy_*) genetic divergence between the two cytotypes for DEGs and NDEs. We observed that DEGs exhibited significantly higher *d_xy_* compared to NDEs, while no significant differences were observed for *F_ST_*. These findings suggest that genes differentially expressed between diploids and tetraploids have accumulated a significantly higher average number of nucleotide differences between ploidies, while nucleotide diversity within each ploidy type remained relatively stable (Fig. 5a,b). Additionally, we compared the ratio of nucleotide diversity at 0-fold to 4-fold degenerate sites (π_0fold_/π_4fold_) between DEGs and NDEs and found a substantially increased π_0fold_/π_4fold_ ratio within DEGs. This result aligns with expectations of a higher mutation load and reduced purifying selection efficiency for these genes (Fig. 5c). DFE analysis further reinforced these observations, revealing that DEGs showed a significantly lower proportion of novel mutations with likely highly deleterious effects compared to NDEs, implying relaxed purifying selection and reduced functional constraints on these genes (Fig. 5d). Finally, these results were robust to random subsampling of two out of four alleles per site from the genotypes of autotetraploids. In the resampling datasets, *F_ST_* values for DEGs were found to be significantly higher than those for NDEs, although the overall trends remained consistent (Fig. S16-S18).

**Fig. 5.**
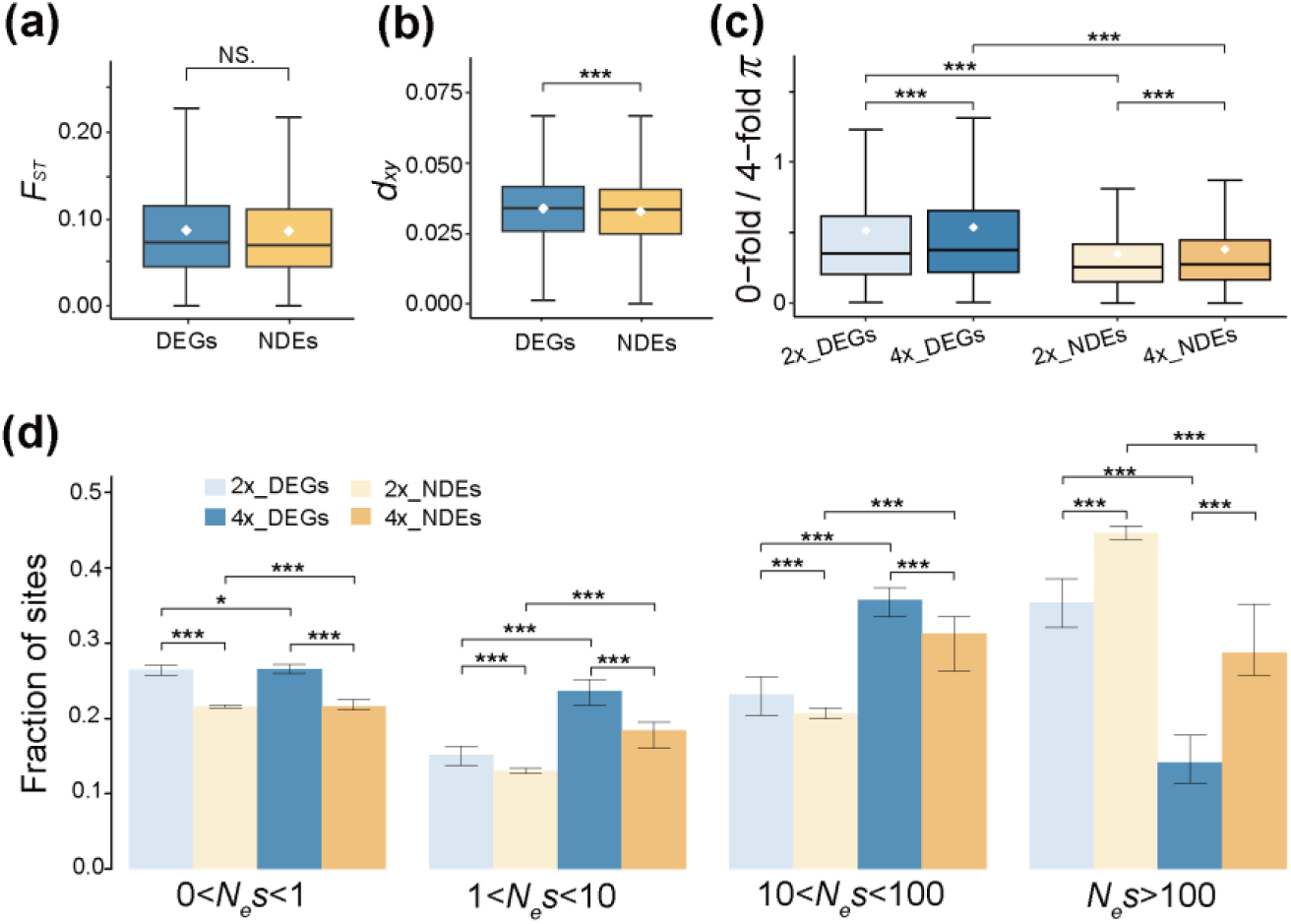
Evolutionary and selection consequences of gene expression changes following autopolyploidization in *O. taibaiensis*. Comparison of genetic divergence of *F_ST_* (a), *d_xy_* (b) between differential expressed genes (DEGs, blue) and non-differentially expressed genes (NDEs, yellow) across samples with different ploidy levels. (c) Comparison of the ratio of 0-fold to 4-fold genetic diversity, used as a measure of selection efficiency, between DEGs and NDEs in diploid (light color) and tetraploid (dark color) populations in *O. taibaiensis*. (d) Distribution of fitness effects (DFE) for DEGs and NDEs in diploid (light color) and tetraploid (dark color) populations in *O. taibaiensis*. Errors bars represent 95% confidence interval based on 200 bootstrap replicates. Asterisks denote significance levels in the Wilcoxon test (two-tailed) (NS. > 0.05, \**P* < 0.05, \*\**P* < 0.01, \*\*\**P* < 0.001).

## 4 Discussion

Polyploidy resulting from WGD is widespread and has repeatedly occurred throughout the evolution of eukaryotes (Masterson 1994; Wood, et al. 2009; Van de Peer, et al. 2017). However, our understanding of its effects on shaping patterns of genomic variation and influencing the selection process remains limited. In this study, we examined the non-model plant genus *Orychophragmus*, which includes both widespread and endemic species, as well as species with variable ploidy levels. By integrating population genomics and transcriptomic datasets, we explored the evolutionary consequences of diverse population demographic histories and the inherent effects of genome doubling on the long-term evolutionary potential of populations.

Using single-copy nuclear genes and population-level SNP datasets, our phylogenomic analyses constructed a more robust species-level phylogeny for this genus compared to previous studies (Hu, et al. 2016; Zhong, et al. 2019). However, we also observed inconsistencies among individual gene trees, likely caused by introgressive hybridization between species (Morales-Briones, et al. 2020; Morales-Cruz, et al. 2021), which merits further investigation in future studies. Notably, *O. taibaiensis* is a locally endemic mountainous species that occupies the highest elevations within the genus and is the only known species to exhibit a mixed-ploidy system. Despite this, little is known about the spatial distribution of different cytotypes across landscapes or the establishment history of polyploids (Barker, et al. 2016). To address this, we first examined the divergence history of *O. taibaiensis* and the closely related, predominantly widespread species *O. violaceus* within the genus. Ecological modeling and population genomic analyses revealed that the two species have experienced divergent demographic histories following their divergence. Specifically, *O. taibaiensis* underwent a more drastic population contraction and a mild recent population size recovery, whereas *O. violaceus* maintained a more stable demographic history. These divergent demographic trajectories have resulted in differences in selection efficacy between the two species. The lower effective population size of *O. taibaiensis* has likely led to the accumulation of a greater deleterious mutation load due to reduced selection efficacy in this species (Harris and Nielsen 2016; Liu, et al. 2022). Additionally, we explored signals of divergent selection between the two species and identified genomic regions enriched with genes involved in responses to oxidative stress. These genes are likely crucial for *O. taibaiensis* to survive and thrive in high mountain regions, highlighting the need for further in-depth studies to uncover the adaptive mechanisms of *O. taibaiensis* in such harsh environments.

The diploid-tetraploid variation in *O. taibaiensis* provides a unique study system to investigate the origin of polyploids, the niche similarity and differentiation of ploidies across spatial scales, and the genomic and evolutionary consequences of genome doubling (Parisod, et al. 2010; Padilla-García, et al. 2023). For the first time, by integrating extensive field surveys, karyotype analysis, and bioinformatics approaches, our results provide strong evidence supporting the autotetraploid origin model of polyploids. Additionally, we clearly delineate the geographic distribution of the two cytotypes across spatial scales. Despite being generally separated by a mountain, we found little environmental differentiation between the cytotypes at local spatial scales across the distribution range of *O. taibaiensis*. Population demographic analyses further revealed that the two cytotypes diverged approximately 328 KYA, with close phylogenetic relationships among individuals of both cytotypes further supporting the relatively recent autopolyploid origin of *O. taibaiensis*.

To better understand how autotetraploids evolve and are affected by selection processes, we examined and compared the distribution of fitness effects of new mutations and the genetic load, estimated as the ratio of 0-fold to 4-fold diversity. Our findings suggest that polysomic masking in autotetraploids, compared to diploids, likely reduces the efficacy of purifying selection. As a result, this relaxed purifying selection leads to a higher accumulation of deleterious mutations in tetraploid populations (Cheng, et al. 2018; Monnahan, et al. 2019). In the short term, the masking of deleterious mutations and the relaxed purifying selection that follow genome doubling may facilitate the rapid establishment and niche expansion of autotetraploid populations (Van de Peer, et al. 2017; Baduel, et al. 2019). However, it remains uncertain how the accumulation of deleterious mutations will balance against the beneficial effects of masking in the long run, and how this dynamic interplay will shape the persistence and coexistence of the two cytotypes.

Moreover, nuclear volume changes arising from polyploidization can increase genome complexity and influence gene expression (Burns, et al. 2021; Yu, et al. 2023). By analyzing gene expression patterns in leaf and root tissues of the two cytotypes of *O. taibaiensis*, we identified a set of differentially expressed genes (DEGs) between diploids and autotetraploids, while the majority of genes were not differentially expressed (NDEs). Consistent with the expected nuclear volume changes, DEGs were significantly enriched in functions related to extracellular structure organization, cell wall assembly, and cell communication (Westermann 2021; Zhu, et al. 2024). Additionally, genes involved in signal transduction, response to stimuli, defense responses, regulation of primary and secondary metabolic processes, and circadian rhythm were also enriched among DEGs. These findings provide a likely explanation for the frequent association of polyploidization with enhanced stress resistance and local environmental adaptation following divergence from diploid relatives (Parisod, et al. 2010; McDaniel 2024). Strikingly, we found that genes with expression changes between ploidy cytotypes evolve under relaxed selective constraints and accumulate more sequence divergence compared to NDEs. This highlights a strong association between expression and sequence changes in response to genome doubling for these genes. Future studies are needed to uncover the potential evolutionary forces driving this association and to explore how selection and transcriptional mechanisms jointly respond to genome doubling, shaping the subsequent evolution of autopolyploids (Bhaskara, et al. 2023; Louder, et al. 2024).

Importantly, there are a few caveats that must be acknowledged. First, biases may arise because many populations genetic analyses assume a diploid model of allele frequencies at mutation-selection-drift balance, which could affect and bias the estimates for autotetraploids (Monnahan, et al. 2019; Yu, et al. 2023). In this study, we called variants in both diploid and polyploid modes for the autotetraploids and also performed random subsampling of two alleles per site to facilitate comparisons across various datasets. While we found consistent results, we must acknowledge that we cannot completely rule out the possibility of bias. Second, our transcriptome analyses compared the relative expression levels of genes to the total transcriptome between the two cytotypes. However, we did not estimate the true transcriptome sizes due to the lack of normalization of transcripts for cell number and biomass content changes in autotetraploids (Coate 2023; Srikant, et al. 2024). This means we cannot entirely exclude the possibility that an overall change in the total number of transcripts occurred following genome doubling. Nevertheless, given that the majority of genes showed balanced expression levels between cytotypes, we believe the likelihood of substantial changes in cell number and transcriptome sizes in autotetraploids is low.

Altogether, our findings provide a novel empirical study system to explore the genomic consequences of genome doubling and the potential evolutionary drivers underlying the successful establishment of newly formed autotetraploid lineages in the local mountainous endemic species *O. taibaiensis*. Future work could focus on sampling and sequencing a broader range of diploids and autotetraploids to better understand the factors influencing the evolutionary potential and establishment of polyploids, as well as their long-term coexistence with their diploid ancestors.

## Acknowledgments

This work was supported by the National Natural Science Foundation of China (32000265) and Fundamental Research Funds for the Central Universities (2023SCUNL105) to J.W.

## Author contributions

J.W. conceived and supervised the study. Q.L., C.J., R.W., Y.L., Y.H. handled the sampling, material collection and performed experiments. Q.L., Z.W., X. Q., Z. Z. analyzed the data, Q.L. and J.W. wrote the manuscript with the input from P.K.I and J.L.. All authors approved the final version of the manuscript.

## Competing interests

The authors declare no competing interests.

## Data availability

All data needed to evaluate the conclusions in this study are present in the paper and/or the Supplementary information. The newly generated whole-genome resequencing data and transcriptome data of the samples produced in this study have been deposited in the National Genomics Data Center (https://ngdc.cncb.ac.cn) under the accession number PRJCA035915.

## Notes

https://ngdc.cncb.ac.cn/search/all?&q=PRJCA035915

### Competing Interest Statement

The authors have declared no competing interest.

### Summary of Updates

The subject area of the manuscript has been updated from molecular biology to evolutionary biology. The Latin names of species in the abstract and title have been corrected to standard italic formatting.

