## supplemental information, Table S1-S9; Fig. S1-S18 for "Evolutionary responses and genomic consequences of polyploidization in natural populations of *Orychophragmus*"

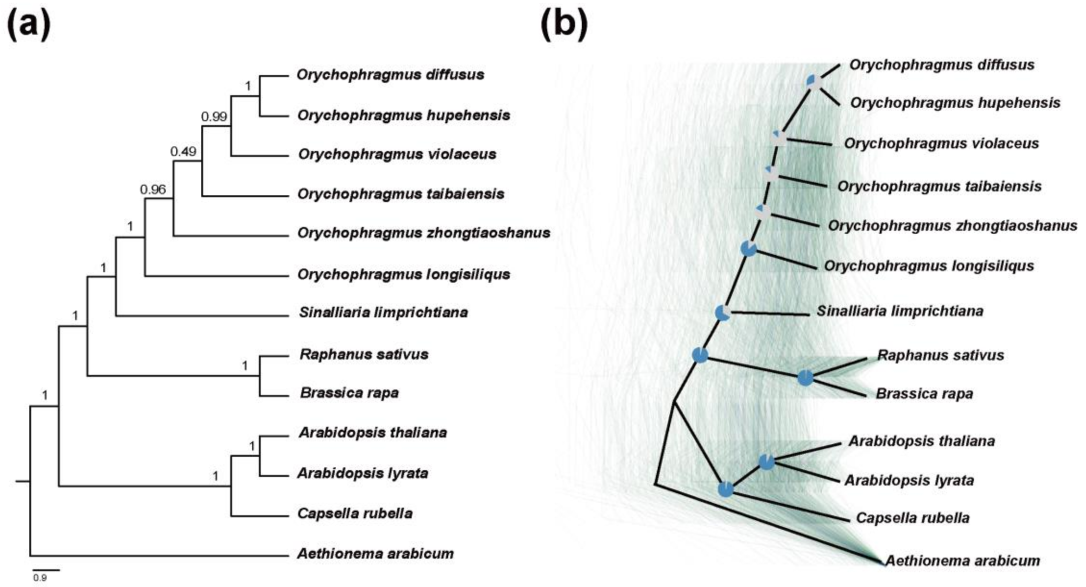


**Fig. S1** Phylogenetic relationships among six *Orychophragmus* species and its relative species based on 514 single-copy genes. (a) Phylogenetic relationships of *Orychophragmus* species inferred using the Astral coalescent method, with branch values indicating bootstrap support. (b) Visualization of 514 gene trees using DensiTree, where thin lines represent individual gene trees, and the thick black line represents the species tree. Pie charts on each branch illustrate the proportion of gene trees supporting the branch structure (blue) versus those not supporting it (gray).

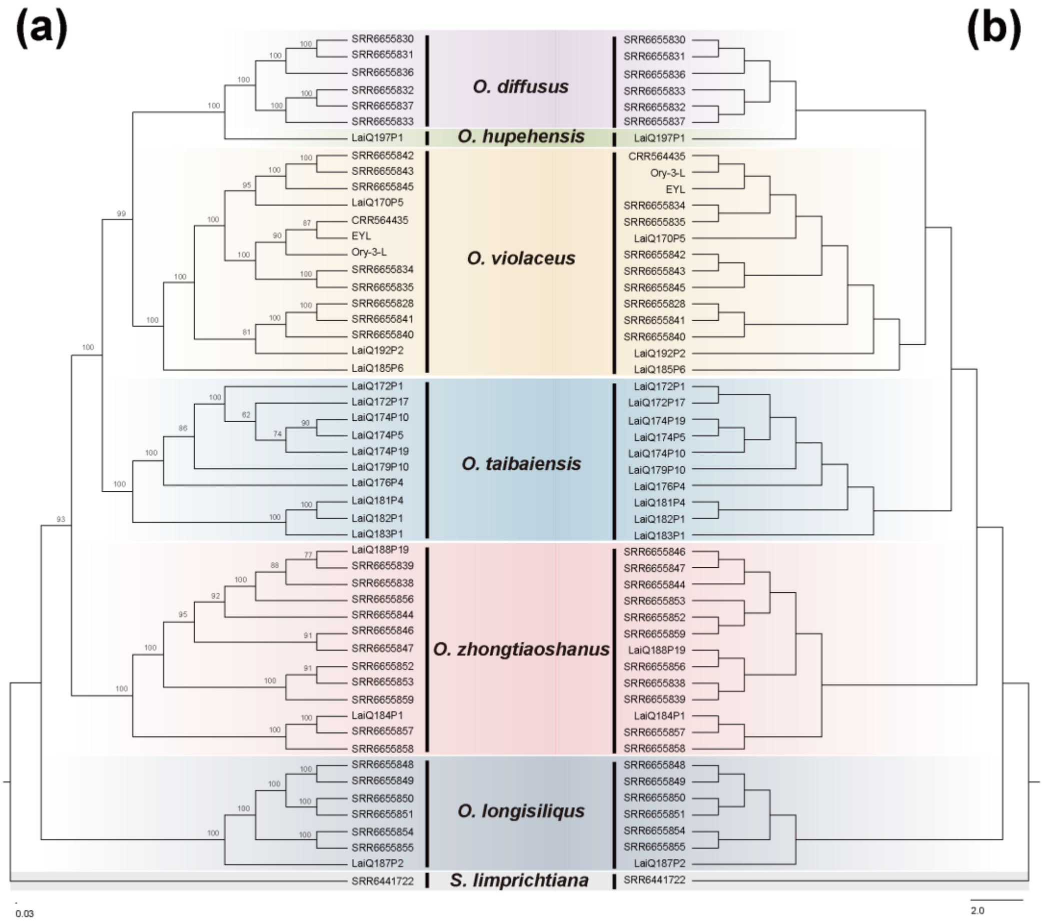


**Fig. S2** Phylogenetic relationships of *Orychophragmus* based on population SNPs data. (a) Phylogenetic tree constructed using the maximum likelihood (ML) method, with branch values indicating bootstrap support. (b) Phylogenetic tree constructed using the neighbor-joining (NJ) method.

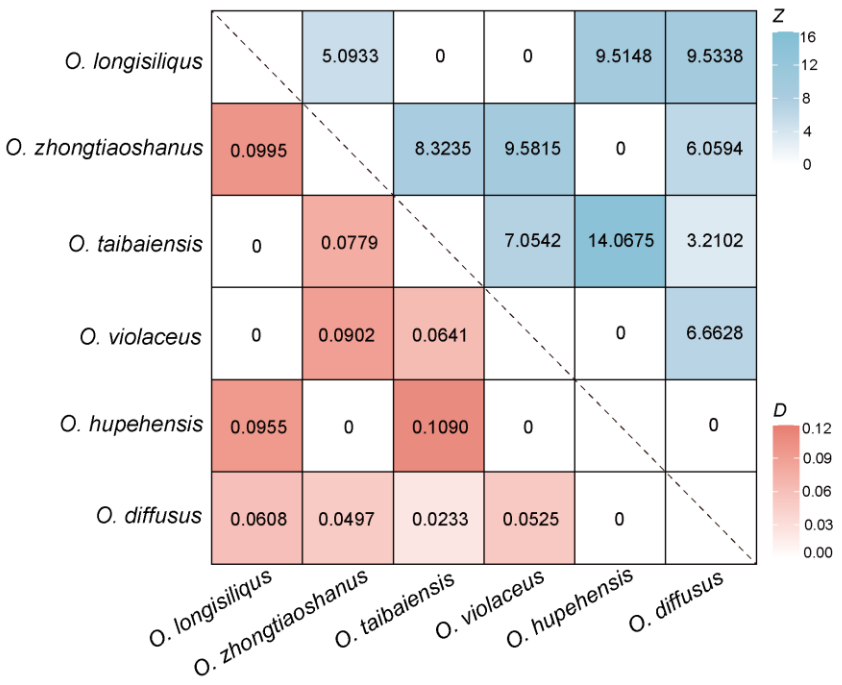


**Fig. S3** Heatmap indicating the maximum pairwise *D* statistics (below diagonal) and the corresponding *Z*-scores (above diagonal) between species pairs across all combinations of trios estimated using Dsuite.

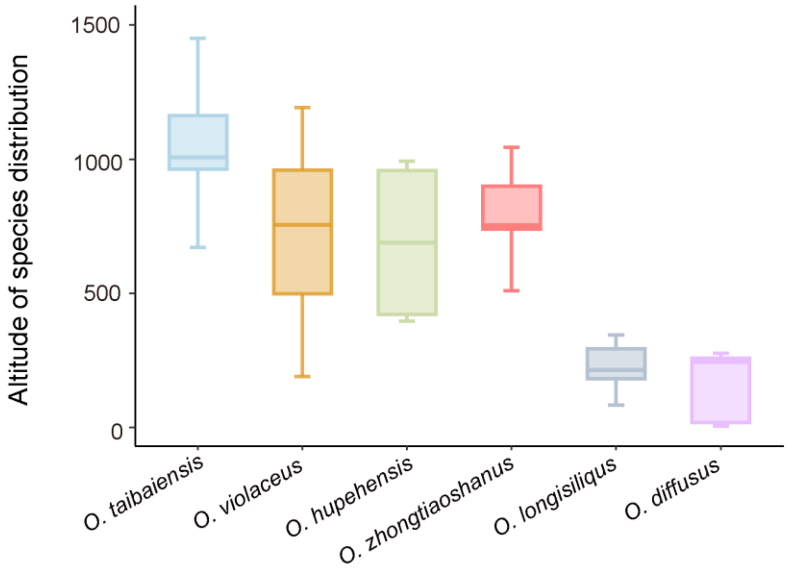


**Fig. S4** The elevation distribution of the six *Orychophragmus* species was compared based on field investigations conducted in this study, records from the Chinese Virtual Herbarium (CVH, https://www.cvh.ac.cn/), and previous research (Hu et al., 2015).

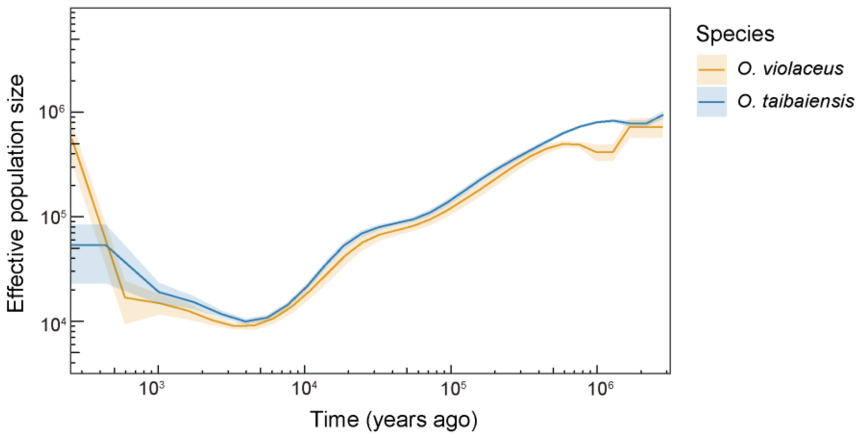


**Fig. S5** The historical effective population size of the *O. violaceus* and *O. taibaiensis* was inferred using MSMC2 based on sets of eight haplotypes for each species. Solid lines representing medians and shading representing ± standard deviation calculated across pairs of haplotypes.

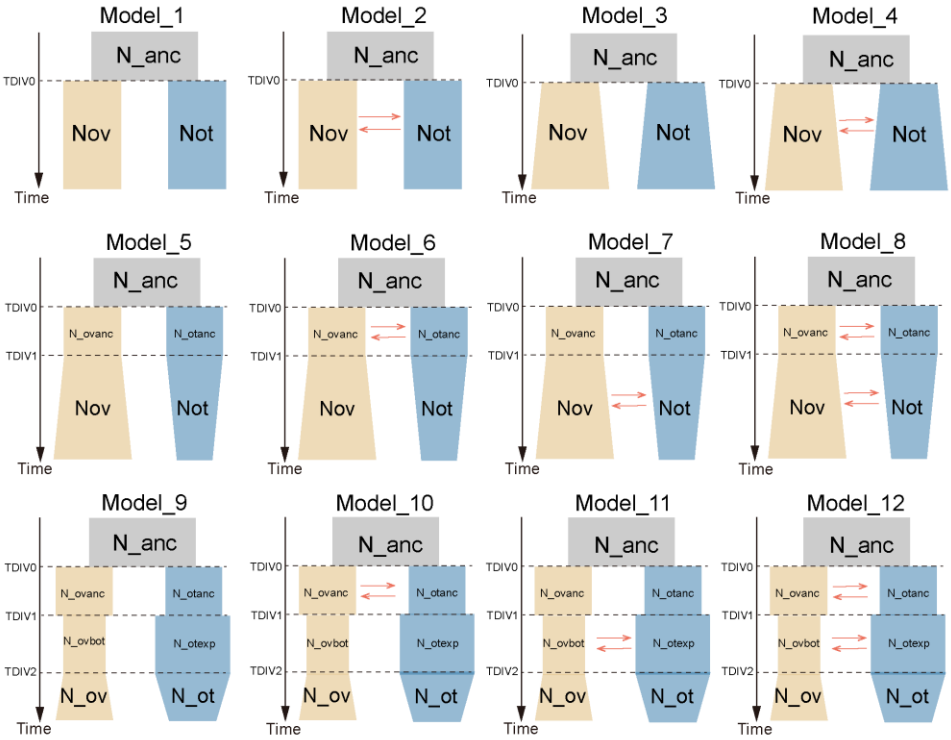


**Fig. S6** Twelve tested demographic models for fastsimcoal2 simulations. Four isolation-without-gene-flow (Model 1, 3, 5, and 9) and eight isolation with-gene-flow models (Model 2, 4, 6, 7,8,10,11and 12). In models 1 and 2, descendant populations exhibited consistent population sizes. Models 3 and 4 showed expansion immediately after splitting. Models 5 to 8 involved expansion of *O. violaceus* but a bottleneck of *O. taibaiensis* at time TDIV1. Models 9 to 12 featured a bottleneck of *O. violaceus* but expansion of *O. taibaiensis* at time TDIV1, followed by expansion of *O. violaceus* and a bottleneck of *O. taibaiensis* at time TDIV2.

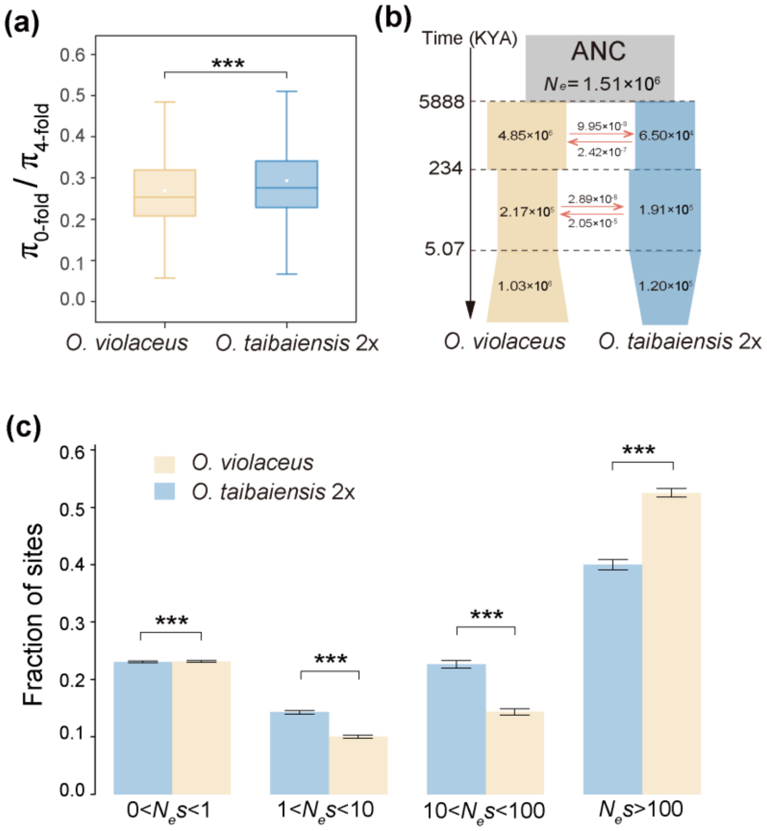


**Fig. S7** Estimation of population divergence history and purifying selection efficiency between *O. violaceus* (yellow) and diploid *O. taibaiensis* (blue). (a) The ratio of nucleotide diversity at 0-fold sites relative to 4-fold sites. (b) The best-fit divergence model inferred by *fastsimcoal2*. Each block represents a current or ancestral population, with arrows denoting ongoing gene flow after divergence (per generation migration rate). The timing of historical events is shown in kilo years ago (KYA). (c) Distribution of fitness effects (DFE) in bins of $N_e$s for new 0-fold nonsynonymous mutations for *O. violaceus* and diploid *O. taibaiensis*. Errors bars represent 95% confidence interval based on 200 bootstrap replicates. Asterisks above the boxplot indicate the level of significance in Wilcoxon test (two-tailed) (****P* < 0.001).

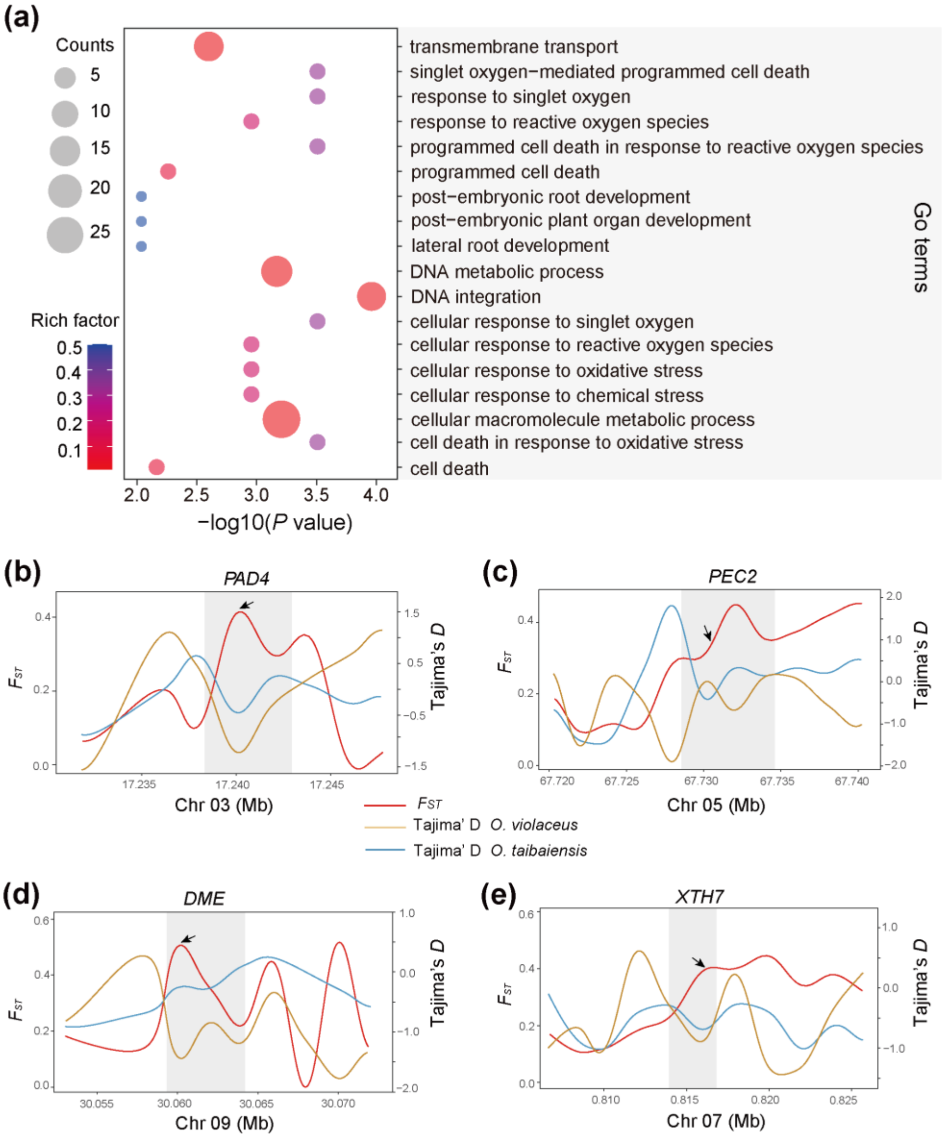


**Fig. S8** Divergent selection detection between *O. violaceus* and *O. taibaiensis* identified using XP-CLR. (a) Gene Ontology (GO) enrichment analysis of genes within candidate selective regions identified by XP-CLR. (b-e) Comparison of $F_{ST}$ and Tajima's D for selected genes within the candidate divergent regions. Shaded areas denote gene regions, while black arrows highlight regions under selection as identified by XP-CLR.

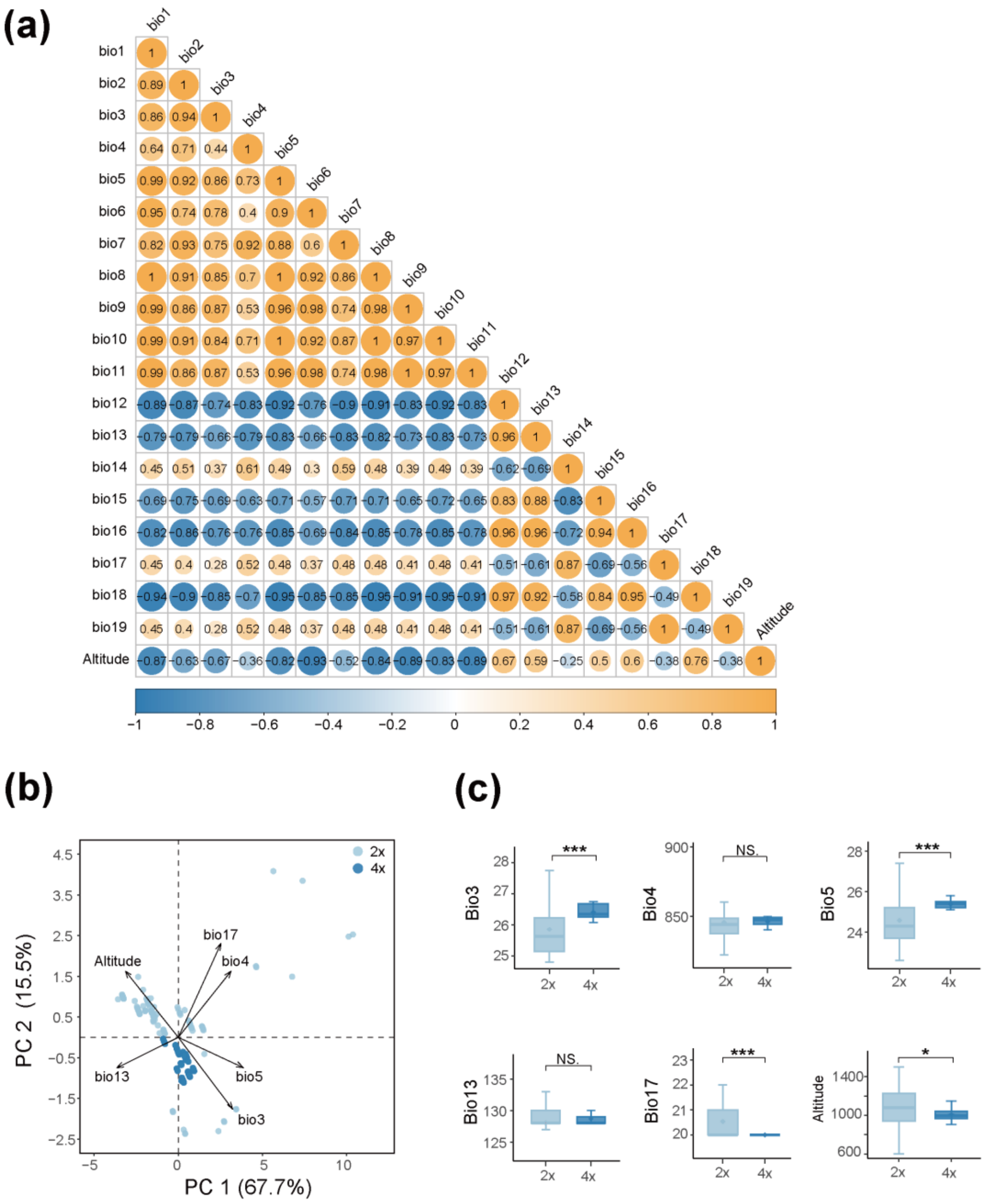


**Fig. S9** Geographical distribution of *O. taibaiensis* at different ploidy levels in relation to climatic differences. (a) Correlation between the 19 environmental factors and altitude within the distribution range of *O. taibaiensis*. (b) PCA plot of *O. taibaiensis* at different ploidy levels, based on five uncorrelated environmental variables and altitude. (c) Comparison of the five environmental variables and altitude for *O. taibaiensis* at different ploidy levels shown in (b).

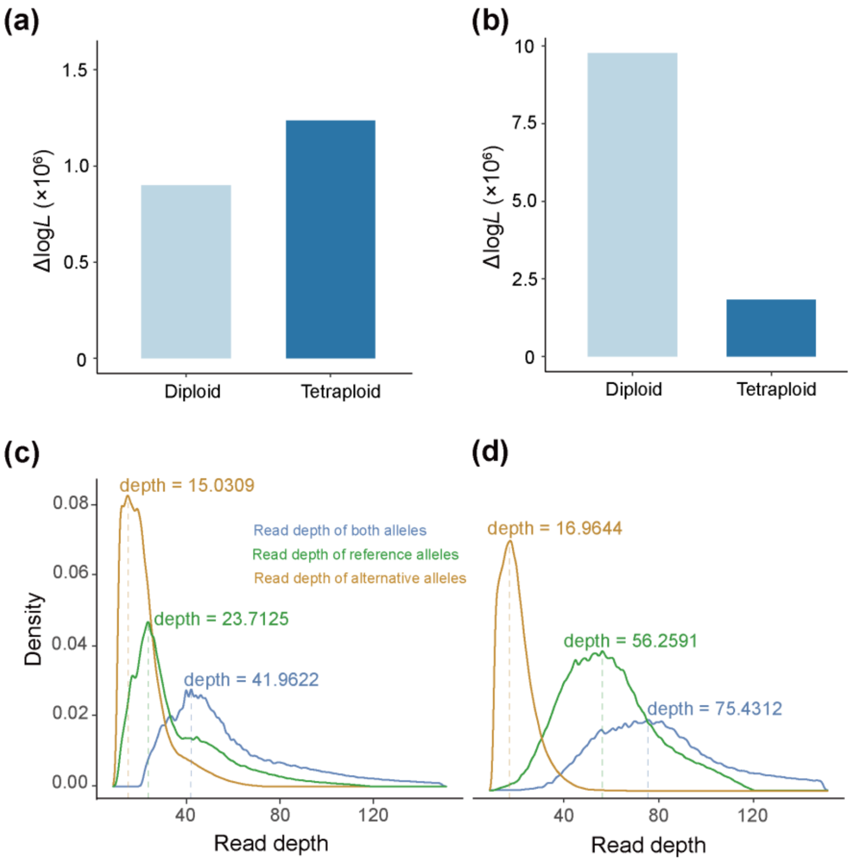


**Fig. S10** Ploidy identification in *O. taibaiensis*. Bar plots show the ΔlogL of all fixed models indicating diploid (a) and tetraploid (b) ploidy levels based on nQuire analysis, where the lowest ΔlogL value corresponds to the ploidy of the individual. (c-d) Read depth density for reference, alternative, and combined alleles in *O. taibaiensis*. In diploids (c), the read depths of alternative and reference alleles are similar, whereas in tetraploids (d), the read depth of the reference alleles is one-third that of the alternative alleles.

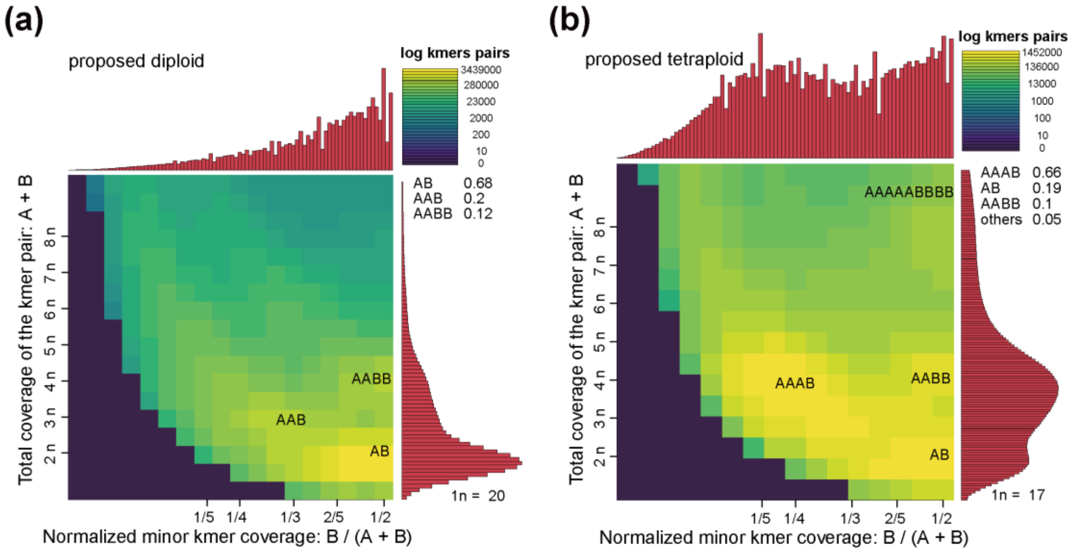


**Fig. S11** Smudgeplots of different ploidy levels in *O. taibaiensis*. (a) Smudgeplots for the diploid *O. taibaiensis*. (b) Smudgeplots for the tetraploid *O. taibaiensis*.

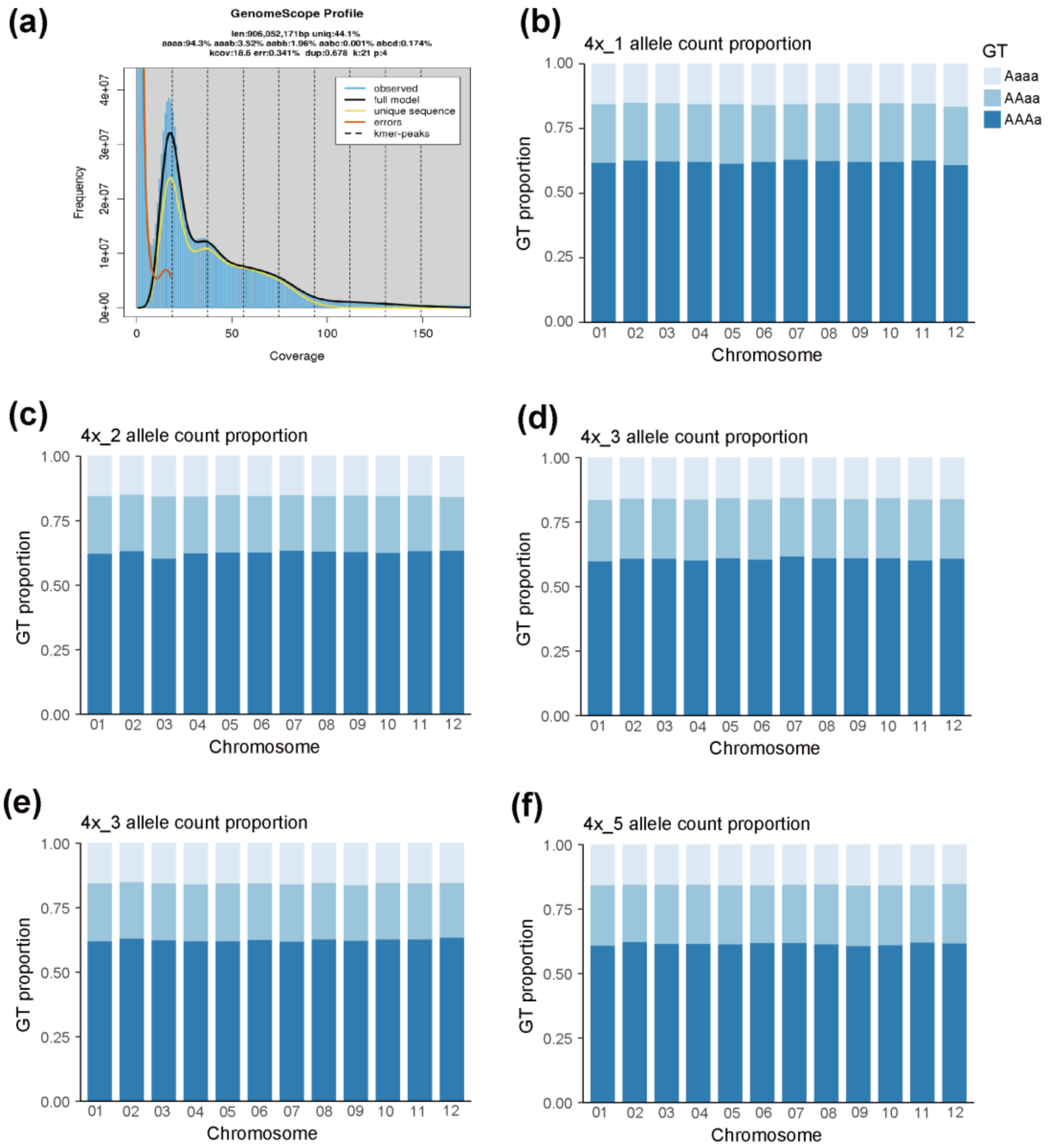


**Fig. S12** (a) *K-mer* spectra and fitted models for the tetraploid individual, showing nucleotide heterozygosity patterns aaab (3.52%) > aabb (1.96%), indicating that the tetraploid is an autotetraploid. (b-f) The relative proportions of various genotypes along the twelve chromosomes for each of the five tetraploid *O. taibaiensis* individuals.

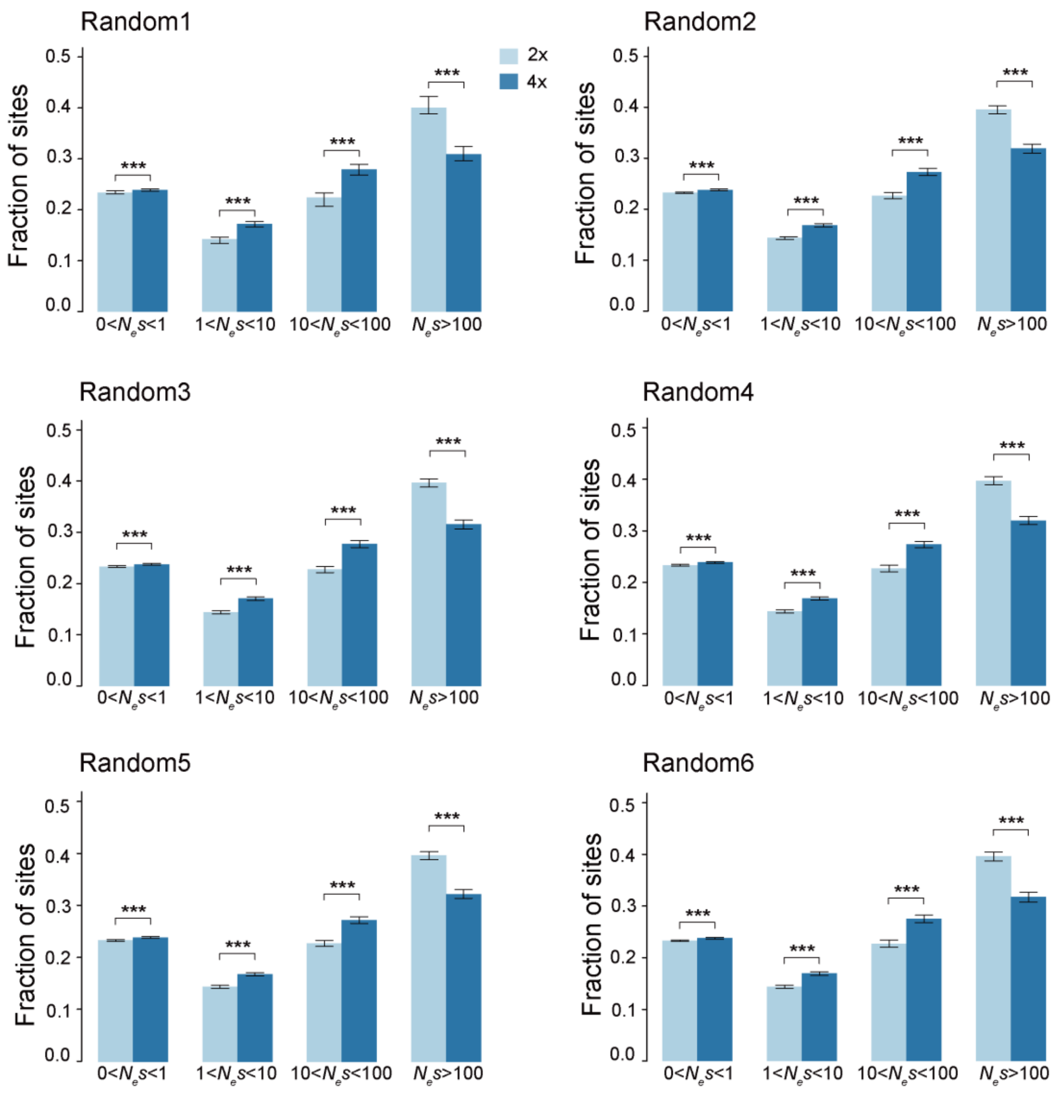


**Fig. S13** Comparison of distribution of fitness effects (DFE) in bins of $N_e$s for new 0-fold nonsynonymous mutations between diploid and tetraploid *O. taibaiensis* based on six random subsampling datasets. For the tetraploid genotypes, two alleles were randomly subsampled from the four alleles per site to generate the random subsampling datasets. Errors bars represent 95% confidence interval based on 200 bootstrap replicates. Asterisks denote significance levels in the Wilcoxon test (two-tailed) (NS. > 0.05, **P* < 0.05, ***P* < 0.01, ****P* < 0.001).

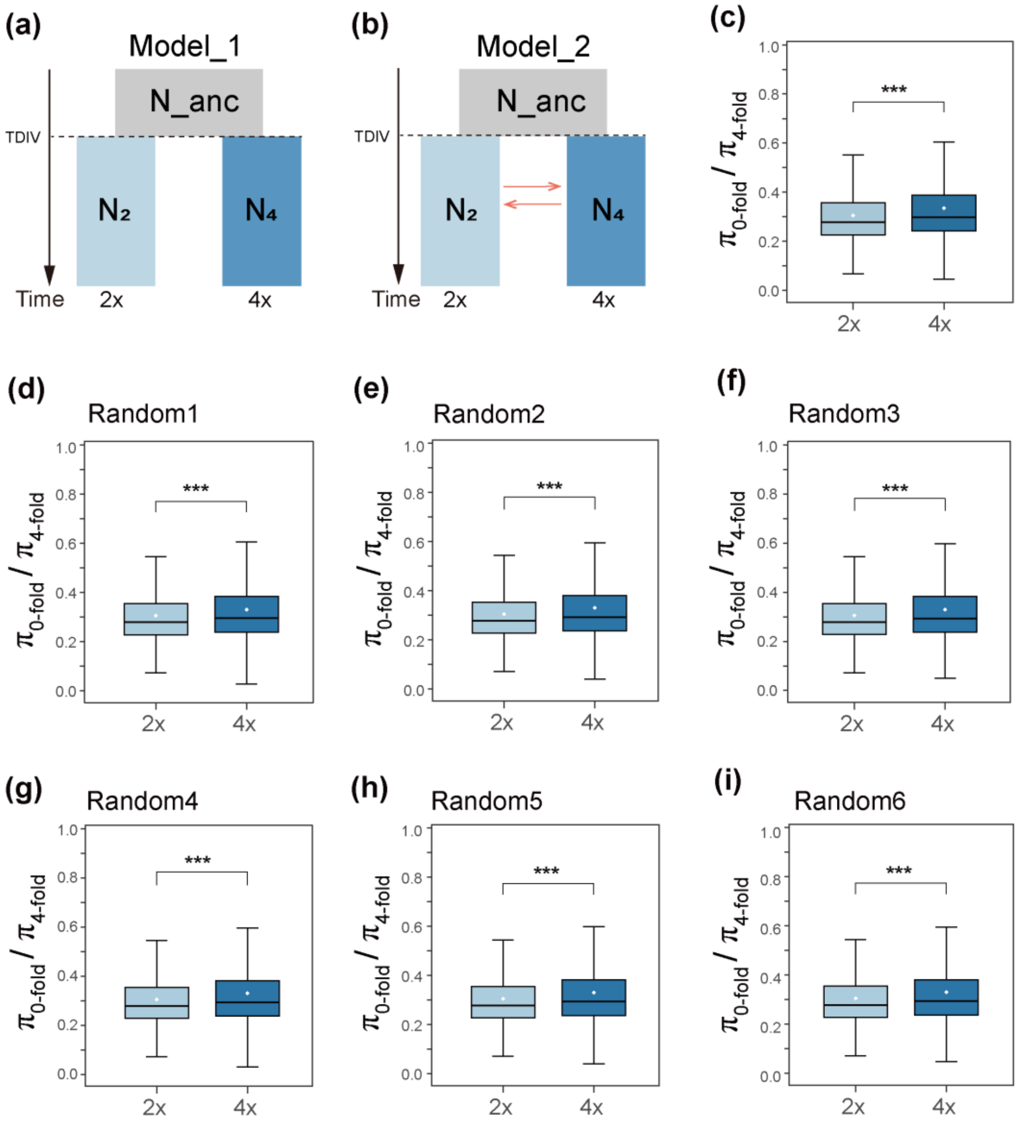


**Fig. S14** Demographic models without (a) and with (b) gene flow inferred for diploid and autotetraploid O. taibaiensis using fastsimcoal2. (c-i) Comparison of the ratio of nucleotide diversity at 0-fold to 4-fold sites between diploid and autotetraploid O. taibaiensis in real data (c), along with the results based on six random subsampling datasets (d-i). For the tetraploid genotypes, two alleles were randomly subsampled from the four alleles per site to generate the random subsampling datasets. Asterisks above the boxplot indicate the levels of significance in Wilcoxon test (two-tailed) (***$P$ < 0.001).

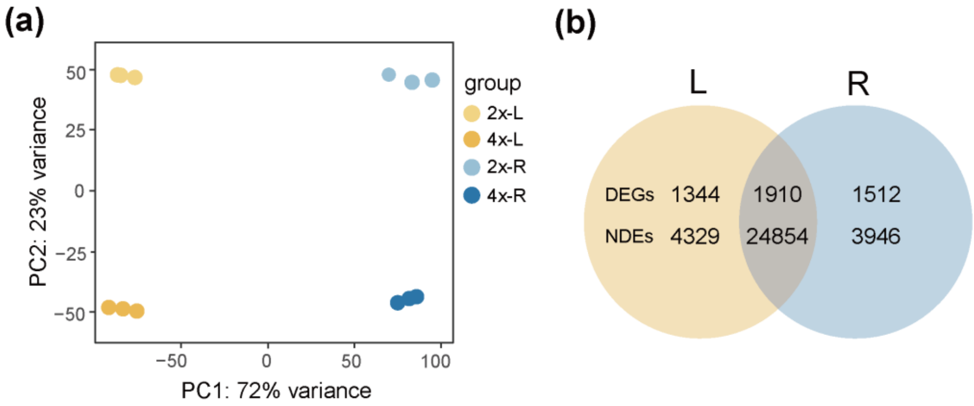


**Fig. S15** Gene expression patterns between diploid and autotetraploid *O. taibaiensis*. (a) Principal component analysis (PCA) of gene expression effectively separates samples by tissue type and ploidy level, with PC1 and PC2 explaining 72% and 23% of the variance, respectively. (b) The number of shared and unique differentially expressed genes (DEGs) and non-differentially expressed genes (NDEs) between diploid and autotetraploid *O. taibaiensis* in leaf (yellow, L) and root tissues (blue, R).

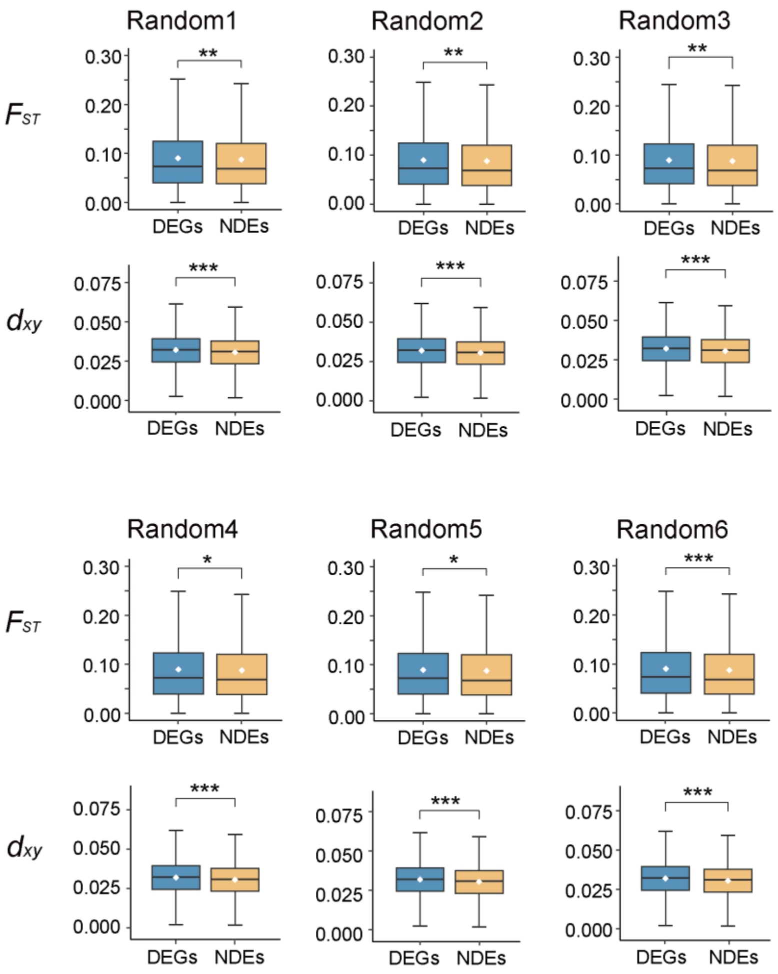


**Fig. S16** Genetic divergence ($F_{ST}$, $d_{xy}$) between diploid and autotetraploid *O. taibaiensis* for differentially expressed genes (DEGs) and non-differentially expressed genes (NDEs) based on six random subsampling datasets. For the tetraploid genotypes, two alleles were randomly subsampled from the four alleles per site to generate the random subsampling datasets. Asterisks indicate significance levels in the Wilcoxon test (two-tailed) (NS. > 0.05, *$P < 0.05$, **$P < 0.01$, ***$P < 0.001$).

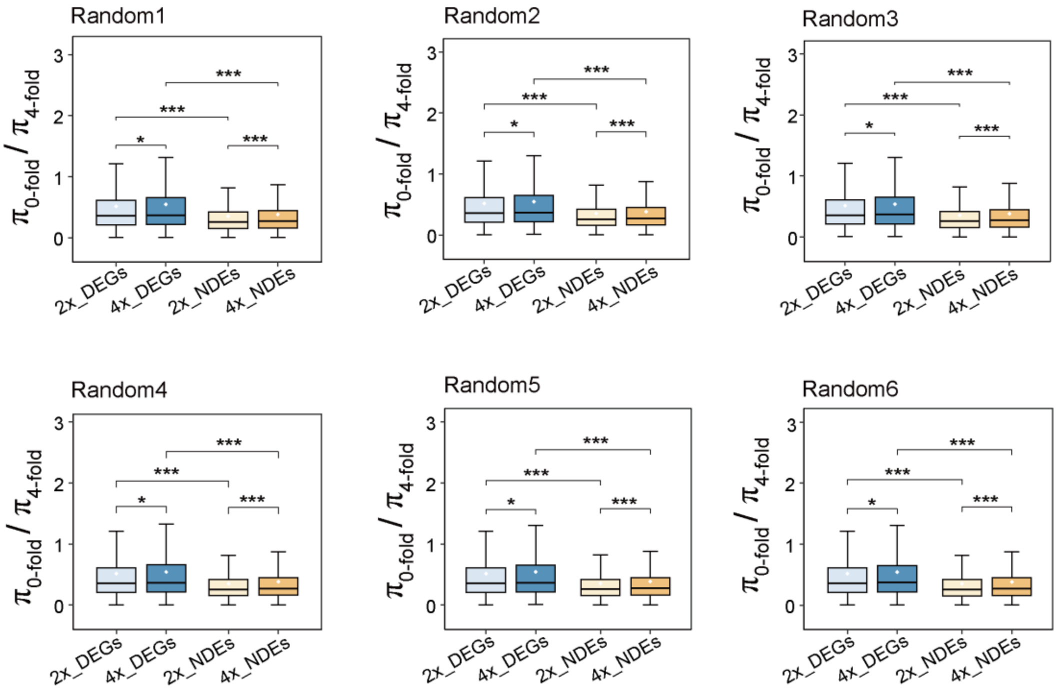


**Fig. S17** Comparison of the ratio of 0-fold to 4-fold genetic diversity, used as a measure of selection efficiency, between differentially expressed genes (DEGs) and non-differentially expressed genes (NDEs) in diploid (light color) and tetraploid (dark color) populations of *O. taibaiensis* based on six random subsampling datasets. For tetraploid genotypes, two alleles were randomly subsampled from the four alleles per site to generate the random subsampling datasets. Asterisks indicate significance levels from the Wilcoxon test (two-tailed): NS (not significant) > 0.05, P < 0.05, *P < 0.01, **P < 0.001.

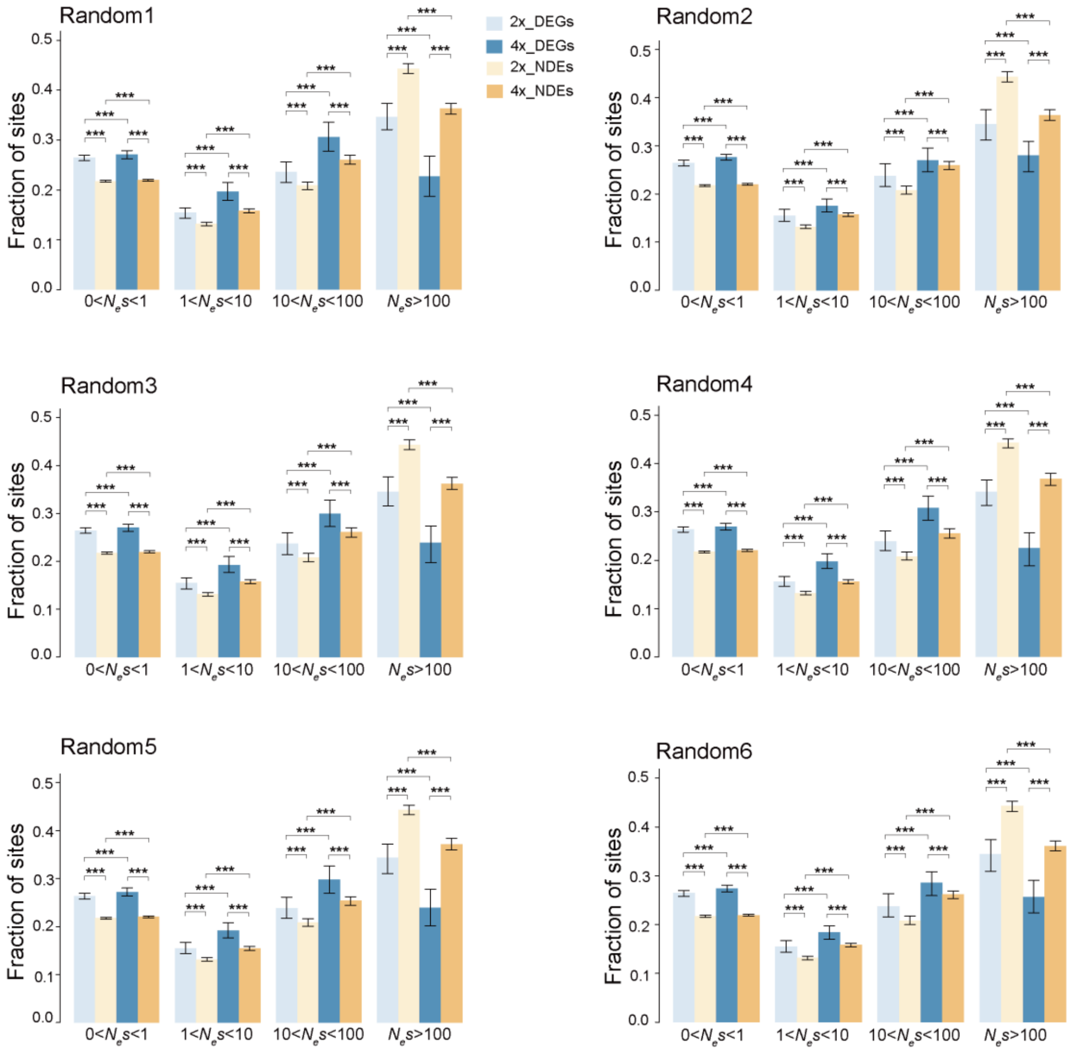


**Fig. S18** Distribution of fitness effects (DFE) for differentially expressed genes (DEGs) and non-differentially expressed genes (NDEs) in diploid (light color) and tetraploid (dark color) populations of *O. taibaiensis* based on six random subsampling datasets. For tetraploid genotypes, two alleles were randomly subsampled from the four alleles per site to generate the random subsampling datasets. Errors bars represent 95% confidence interval based on 200 bootstrap replicates. Asterisks indicate significance levels in the Wilcoxon test (two-tailed) (NS. > 0.05, *$P < 0.05$, **$P < 0.01$, ***$P < 0.001$).

**Table S1** Summary of data sources used for phylogenetic and population analyses, along with sequencing information.

| Species | Data type | Sample ID | Coverage ratio | Mapped ratio | Depth | Sources |
|---|---|---|---|---|---|---|
| *O. taibaiensis* | Resequencing | LaiQ172P1/4x_1 | 0.840543 | 0.9629 | 65.2004 | This study |
| *O. taibaiensis* | Resequencing | LaiQ172P17/4x_2 | 0.855916 | 0.9595 | 80.5837 | This study |
| *O. taibaiensis* | Resequencing | LaiQ174P10/4x_3 | 0.809907 | 0.9675 | 51.8769 | This study |
| *O. taibaiensis* | Resequencing | LaiQ174P19/4x_4 | 0.850027 | 0.9547 | 82.4459 | This study |
| *O. taibaiensis* | Resequencing | LaiQ174P5/4x_5 | 0.822669 | 0.976 | 56.386 | This study |
| *O. taibaiensis* | Resequencing | LaiQ176P4/2x_1 | 0.784046 | 0.9653 | 50.8453 | This study |
| *O. taibaiensis* | Resequencing | LaiQ179P10/2x_2 | 0.79656 | 0.9591 | 53.359 | This study |
| *O. taibaiensis* | Resequencing | LaiQ181P4/2x_3 | 0.784309 | 0.956 | 44.2196 | This study |
| *O. taibaiensis* | Resequencing | LaiQ182P1/2x_4 | 0.716503 | 0.9661 | 20.3266 | This study |
| *O. taibaiensis* | Resequencing | LaiQ183P1/2x_5 | 0.760726 | 0.9643 | 19.6349 | This study |
| *O. violaceus* | Resequencing | LaiQ170P5 | 0.932832 | 0.9829 | 39.9179 | This study |
| *O. violaceus* | Resequencing | LaiQ185P6 | 0.89747 | 0.9489 | 54.7158 | This study |
| *O. violaceus* | Resequencing | LaiQ192P2 | 0.91156 | 0.981 | 42.9088 | This study |
| *O. violaceus* | Resequencing | CRR564435 | 0.941259 | 0.9866 | 59.5594 | Huang, et al. 2023 |
| *O. violaceus* | Resequencing | EYL | 0.911188 | 0.9788 | 63.6742 | Zhang, et al. 2023 |
| *O. violaceus* | Resequencing | Ory-3-L | 0.999249 | 0.9921 | 48.5668 | Jia, et al. 2023 |
| *O. violaceus* | RNAseq | SRR6655828 | - | - | - | Zhong, et al. 2019 |
| *O. violaceus* | RNAseq | SRR6655834 | - | - | - | Zhong, et al. 2019 |
| *O. violaceus* | RNAseq | SRR6655835 | - | - | - | Zhong, et al. 2019 |
| *O. violaceus* | RNAseq | SRR6655840 | - | - | - | Zhong, et al. 2019 |
| *O. violaceus* | RNAseq | SRR6655841 | - | - | - | Zhong, et al. 2019 |
| *O. violaceus* | RNAseq | SRR6655842 | - | - | - | Zhong, et al. 2019 |
| *O. violaceus* | RNAseq | SRR6655843 | - | - | - | Zhong, et al. 2019 |
| *O. violaceus* | RNAseq | SRR6655845 | - | - | - | Zhong, et al. 2019 |

Continued Table S1

| Species | Data type | Sample ID | Coverage ratio | Mapped ratio | Depth | Sources |
|---|---|---|---|---|---|---|
| *O. longisiliqus* | Resequencing | LaiQ187P2 | 0.702945 | 0.9529 | 43.6717 | This study |
| *O. longisiliqus* | RNAseq | SRR6655848 | - | - | - | Zhong, et al. 2019 |
| *O. longisiliqus* | RNAseq | SRR6655849 | - | - | - | Zhong, et al. 2019 |
| *O. longisiliqus* | RNAseq | SRR6655850 | - | - | - | Zhong, et al. 2019 |
| *O. longisiliqus* | RNAseq | SRR6655851 | - | - | - | Zhong, et al. 2019 |
| *O. longisiliqus* | RNAseq | SRR6655854 | - | - | - | Zhong, et al. 2019 |
| *O. longisiliqus* | RNAseq | SRR6655855 | - | - | - | Zhong, et al. 2019 |
| *O. zhongtiaoshanus* | Resequencing | LaiQ184P1 | 0.747125 | 0.9558 | 46.8094 | This study |
| *O. zhongtiaoshanus* | Resequencing | LaiQ188P19 | 0.745104 | 0.9434 | 41.2263 | This study |
| *O. zhongtiaoshanus* | RNAseq | SRR6655838 | - | - | - | Zhong, et al. 2019 |
| *O. zhongtiaoshanus* | RNAseq | SRR6655839 | - | - | - | Zhong, et al. 2019 |
| *O. zhongtiaoshanus* | RNAseq | SRR6655844 | - | - | - | Zhong, et al. 2019 |
| *O. zhongtiaoshanus* | RNAseq | SRR6655846 | - | - | - | Zhong, et al. 2019 |
| *O. zhongtiaoshanus* | RNAseq | SRR6655847 | - | - | - | Zhong, et al. 2019 |
| *O. zhongtiaoshanus* | RNAseq | SRR6655852 | - | - | - | Zhong, et al. 2019 |
| *O. zhongtiaoshanus* | RNAseq | SRR6655853 | - | - | - | Zhong, et al. 2019 |
| *O. zhongtiaoshanus* | RNAseq | SRR6655856 | - | - | - | Zhong, et al. 2019 |
| *O. zhongtiaoshanus* | RNAseq | SRR6655857 | - | - | - | Zhong, et al. 2019 |
| *O. zhongtiaoshanus* | RNAseq | SRR6655858 | - | - | - | Zhong, et al. 2019 |
| *O. zhongtiaoshanus* | RNAseq | SRR6655859 | - | - | - | Zhong, et al. 2019 |
| *O. hupehensis* | Resequencing | LaiQ197P1 | 0.751662 | 0.9593 | 49.547 | This study |
| *O. diffusus* | RNAseq | SRR6655830 | - | - | - | Zhong, et al. 2019 |
| *O. diffusus* | RNAseq | SRR6655831 | - | - | - | Zhong, et al. 2019 |
| *O. diffusus* | RNAseq | SRR6655832 | - | - | - | Zhong, et al. 2019 |

Continued Table S1

| Species | Data type | Sample ID | Coverage ratio | Mapped ratio | Depth | Sources |
|---|---|---|---|---|---|---|
| *O. diffusus* | RNAseq | SRR6655833 | - | - | - | Zhong, et al. 2019 |
| *O. diffusus* | RNAseq | SRR6655836 | - | - | - | Zhong, et al. 2019 |
| *O. diffusus* | RNAseq | SRR6655837 | - | - | - | Zhong, et al. 2019 |
| *S. limprichtiana* | RNAseq | SRR6441722 | - | - | - | Zhong, et al. 2019 |

157

**Table S2** The relative likelihood and AIC values of the 12 models listed in fastsimcoal2 simulation Fig. S6.

| Models | Δlikelihood | AIC |
|---|---|---|
| Model_1 | 45801.939 | 8536993.968 |
| Model_2 | 9983.039 | 8372045.838 |
| Model_3 | 45042.732 | 8533503.691 |
| Model_4 | 9929.839 | 8371806.843 |
| Model_5 | 60104.492 | 8602865.659 |
| Model_6 | 45237.979 | 8534406.836 |
| Model_7 | 296540.889 | 9691697.505 |
| Model_8 | 37856.086 | 8500415.963 |
| Model_9 | 44578.723 | 8531372.85 |
| Model_10 | 10041.963 | 8372329.193 |
| Model_11 | 9956.788 | 8371936.947 |
| **Model_12** | **9837.194** | **8371390.197** |

Note: The smallest Δlikelihood and AIC indicate the best-fit model.

**Table S3** The point estimates and 95% confidence intervals of parameters in the best-fit model (model 12) inferred by fastsimcoal2.

| Parameters | Point estimate | 2.5% Percentile | 97.5% Percentile |
|---|---|---|---|
| N_anc | 1863339 | 672538.95 | 5696666.4 |
| N_ovanc | 778548 | 484639.975 | 888925.7 |
| N_otanc | 60455 | 42330.05 | 73897.425 |
| N_ovbot | 197139 | 192462.225 | 233766.3 |
| N_otexp | 197701 | 193329.6 | 235558.35 |
| Nov | 1083997 | 845056.95 | 1293472.625 |
| Not | 159493 | 145050 | 199365.275 |
| TDIV0 | 5245423 | 1391004.675 | 7326630.6 |
| TDIV1 | 214649 | 205249.425 | 260880.425 |
| TDIV2 | 5215 | 5015.85 | 5788.525 |
| migr21 | 2.458E-05 | 1.629E-05 | 2.458E-05 |
| migr12 | 2.405E-06 | 2.078E-06 | 3.573E-06 |
| migr34 | 1.703E-09 | 3.342E-10 | 2.265E-07 |
| migr43 | 1.147E-07 | 1.228E-09 | 5.015E-06 |
| GR_ov | -3.268E-04 | -3.386E-04 | -2.690E-04 |
| GR_ot | 4.118E-05 | 2.331E-05 | 5.906E-05 |
| MaxEstLhood | -1817818.204 | -1817714.21 | -1817263.575 |
| MaxObsLhood | -1807981.01 | -1807981.01 | -1807981.01 |

**Table S4** Gene Ontology (GO) enrichment analysis of genes within candidate selective regions identified by XP-CLR.

| GO.ID | Term | Annotated | Significant | Expected | Fisher.P | Adjust.P | Rich factor |
|---|---|---|---|---|---|---|---|
| GO:0015074 | DNA integration | 840 | 13 | 3.85 | 0.00011 | 0.00093 | 0.015476 |
| GO:0000304 | response to singlet oxygen | 6 | 2 | 0.03 | 0.00031 | 0.00093 | 0.333333 |
| GO:0010343 | singlet oxygen-mediated programmed cell death | 6 | 2 | 0.03 | 0.00031 | 0.00093 | 0.333333 |
| GO:0036473 | cell death in response to oxidative stress | 6 | 2 | 0.03 | 0.00031 | 0.00093 | 0.333333 |
| GO:0071452 | cellular response to singlet oxygen | 6 | 2 | 0.03 | 0.00031 | 0.00093 | 0.333333 |
| GO:0097468 | programmed cell death in response to reactive oxygen species | 6 | 2 | 0.03 | 0.00031 | 0.00093 | 0.333333 |
| GO:0044260 | cellular macromolecule metabolic process | 3165 | 27 | 14.5 | 0.00061 | 0.00151 | 0.008531 |
| GO:0006259 | DNA metabolic process | 1429 | 16 | 6.54 | 0.00067 | 0.00151 | 0.011197 |
| GO:0000302 | response to reactive oxygen species | 11 | 2 | 0.05 | 0.00111 | 0.00167 | 0.181818 |
| GO:0034599 | cellular response to oxidative stress | 11 | 2 | 0.05 | 0.00111 | 0.00167 | 0.181818 |
| GO:0034614 | cellular response to reactive oxygen species | 11 | 2 | 0.05 | 0.00111 | 0.00167 | 0.181818 |
| GO:0062197 | cellular response to chemical stress | 11 | 2 | 0.05 | 0.00111 | 0.00167 | 0.181818 |
| GO:0055085 | transmembrane transport | 1318 | 14 | 6.04 | 0.00249 | 0.00345 | 0.010622 |
| GO:0012501 | programmed cell death | 24 | 2 | 0.11 | 0.00536 | 0.00689 | 0.083333 |
| GO:0008219 | cell death | 27 | 2 | 0.12 | 0.00675 | 0.0081 | 0.074074 |
| GO:0048527 | lateral root development | 2 | 1 | 0.01 | 0.00914 | 0.00914 | 0.5 |
| GO:0048528 | post-embryonic root development | 2 | 1 | 0.01 | 0.00914 | 0.00914 | 0.5 |
| GO:0090696 | post-embryonic plant organ development | 2 | 1 | 0.01 | 0.00914 | 0.00914 | 0.5 |

**Table S5** Environmental variables used in this study derived from WorldClim.

| Code | Variable |
|---|---|
| Temperature-related | |
| Bio1 | Annual mean temperature(℃) |
| Bio2 | Mean diurnal temperature range (℃) |
| Bio3 | Isothermality (BIO2/BIO7) (×100) |
| Bio4 | Temperature seasonality (standard deviation ×100) |
| Bio5 | Maximum temperature of warmest month(℃) |
| Bio6 | Minimum temperature of coldest month(℃) |
| Bio7 | Temperature annual range(℃) |
| Bio8 | Mean temperature of wettest quarter(℃) |
| Bio9 | Mean temperature of driest quarter(℃) |
| Bio10 | Mean temperature of warmest quarter(℃) |
| Bio11 | Mean temperature of coldest quarter(℃) |
| Precipitation-related | |
| Bio12 | Annual precipitation(mm) |
| Bio13 | Precipitation of wettest month(mm) |
| Bio14 | Precipitation of driest month(mm) |
| Bio15 | Precipitation seasonality(mm) |
| Bio16 | Precipitation of wettest quarter(mm) |
| Bio17 | Precipitation of driest quarter (mm) |
| Bio18 | Precipitation of warmest quarter(mm) |
| Bio19 | Precipitation of coldest quarter(mm) |

**Table S6** Environmental variables (1-19) and elevation data for the geographical distribution of diploid and autotetraploid *O. taibaiensis*. (See the attached separate table S6).

**Table S7** The point estimates and 95% confidence intervals of parameters in the best-fit model inferred by fastsimcoal2 for diploid and tetraploid *O. taibaiensis*, and the point estimates from six random sampling data.

| Parameters | Point estimate | 2.5% Percentile | 97.5% Percentile | Random1 | Random2 | Random3 | Random4 | Random5 | Random6 |
|---|---|---|---|---|---|---|---|---|---|
| N2 | 253251 | 240233 | 360891.825 | 283323 | 300998 | 288518 | 296214 | 270999 | 270054 |
| N4 | 152384 | 145369.325 | 205754.4 | 113694 | 115405 | 106425 | 120110 | 123251 | 101371 |
| N_anc | 18575 | 10739.775 | 27068.25 | 27430 | 21849 | 16489 | 16960 | 20445 | 32486 |
| TDIV | 327943 | 325795.225 | 468081.55 | 342810 | 345498 | 325689 | 353947 | 334526 | 306926 |
| migr24 | 1.756E-05 | 1.228E-05 | 1.884E-05 | 1.464E-05 | 1.485E-05 | 1.699E-05 | 1.539E-05 | 1.694E-05 | 1.637E-05 |
| migr42 | 1.448E-05 | 9.919E-06 | 1.484E-05 | 1.183E-05 | 1.134E-05 | 1.301E-05 | 1.078E-05 | 9.480E-06 | 1.368E-05 |
| MaxEstLhood | -1.465E+06 | -1.465E+06 | -1.465E+06 | -1.663E+06 | -1.663E+06 | -1.663E+06 | -1.664E+06 | -1.663E+06 | -1.664E+06 |
| MaxObsLhood | -1.437E+06 | -1.437E+06 | -1.437E+06 | -1.594E+06 | -1.594E+06 | -1.594E+06 | -1.594E+06 | -1.594E+06 | -1.595E+06 |

**Table S8** Transcriptome sequencing data information for differential expression analysis between diploid and tetraploid *O. taibaiensis*.

| Species | Ploidy | Sample ID | Tissue | Clean Reads | Sum Reads/Mb |
|---|---|---|---|---|---|
| *O. taibaiensis* | Diploid | LaiQ219P21-2-L1 | Leaves | 44,827,150 | 6716.76 |
| *O. taibaiensis* | Diploid | LaiQ219P21-2-L2 | Leaves | 45,418,060 | 6805.84 |
| *O. taibaiensis* | Diploid | LaiQ219P21-2-L3 | Leaves | 39,109,924 | 5859.59 |
| *O. taibaiensis* | Diploid | LaiQ219P21-2-R1 | Roots | 46,034,306 | 6898.21 |
| *O. taibaiensis* | Diploid | LaiQ219P21-2-R2 | Roots | 48,402,284 | 7253.75 |
| *O. taibaiensis* | Diploid | LaiQ219P21-2-R3 | Roots | 46,389,480 | 6952.06 |
| *O. taibaiensis* | Tetraploid | LaiQ216P9-1-L1 | Leaves | 44,800,212 | 6713.64 |
| *O. taibaiensis* | Tetraploid | LaiQ216P9-1-L2 | Leaves | 47,723,592 | 7151.83 |
| *O. taibaiensis* | Tetraploid | LaiQ216P9-1-L3 | Leaves | 39,876,634 | 5974.77 |
| *O. taibaiensis* | Tetraploid | LaiQ216P9-1-R1 | Roots | 43,575,798 | 6527.81 |
| *O. taibaiensis* | Tetraploid | LaiQ216P9-1-R2 | Roots | 39,601,462 | 5933.75 |
| *O. taibaiensis* | Tetraploid | LaiQ216P9-1-R3 | Roots | 40,160,344 | 6017.50 |

**Table S9** Summary of Gene Ontology (GO) enrichment terms for differentially expressed genes between diploid and tetraploid *O. taibaiensis*.

| Group | GO.ID | Term | Annotated | Significant | Expected | Fisher.*P* | Adjust.*P* |
|---|---|---|---|---|---|---|---|
| L: 2x up 4x | GO:0007165 | **signal transduction** | 666 | 45 | 17 | 0.00000 | 0.00000 |
| L: 2x up 4x | GO:0023052 | signaling | 667 | 45 | 17.03 | 0.00000 | 0.00000 |
| L: 2x up 4x | GO:0007154 | **cell communication** | 679 | 45 | 17.33 | 0.00000 | 0.00000 |
| L: 2x up 4x | GO:0051716 | cellular response to stimulus | 1113 | 59 | 28.41 | 0.00000 | 0.00000 |
| L: 2x up 4x | GO:0050896 | **response to stimulus** | 1826 | 75 | 46.61 | 0.00002 | 0.00011 |
| L: 2x up 4x | GO:0050789 | **regulation of biological process** | 2888 | 104 | 73.72 | 0.00012 | 0.00048 |
| L: 2x up 4x | GO:0065007 | biological regulation | 3033 | 106 | 77.42 | 0.00032 | 0.00110 |
| L: 2x up 4x | GO:0050794 | regulation of cellular process | 2633 | 93 | 67.21 | 0.00058 | 0.00174 |
| L: 2x up 4x | GO:0031398 | positive regulation of protein ubiquitination | 12 | 3 | 0.31 | 0.00306 | 0.00503 |
| L: 2x up 4x | GO:0051443 | positive regulation of ubiquitin-protein transferase activity | 12 | 3 | 0.31 | 0.00306 | 0.00503 |
| L: 2x up 4x | GO:1903322 | positive regulation of protein modification by small protein conjugation or removal | 12 | 3 | 0.31 | 0.00306 | 0.00503 |
| L: 2x up 4x | GO:1904668 | positive regulation of ubiquitin protein ligase activity | 12 | 3 | 0.31 | 0.00306 | 0.00503 |
| L: 2x up 4x | GO:0031440 | regulation of mRNA 3'-end processing | 4 | 2 | 0.1 | 0.00377 | 0.00503 |
| L: 2x up 4x | GO:0031441 | negative regulation of mRNA 3'-end processing | 4 | 2 | 0.1 | 0.00377 | 0.00503 |
| L: 2x up 4x | GO:0050686 | negative regulation of mRNA processing | 4 | 2 | 0.1 | 0.00377 | 0.00503 |
| L: 2x up 4x | GO:1900363 | regulation of mRNA polyadenylation | 4 | 2 | 0.1 | 0.00377 | 0.00503 |
| L: 2x up 4x | GO:1900364 | negative regulation of mRNA polyadenylation | 4 | 2 | 0.1 | 0.00377 | 0.00503 |
| L: 2x up 4x | GO:1903312 | negative regulation of mRNA metabolic process | 4 | 2 | 0.1 | 0.00377 | 0.00503 |
| L: 2x up 4x | GO:0031396 | regulation of protein ubiquitination | 15 | 3 | 0.38 | 0.00598 | 0.00641 |
| L: 2x up 4x | GO:0051438 | regulation of ubiquitin-protein transferase activity | 15 | 3 | 0.38 | 0.00598 | 0.00641 |
| L: 2x up 4x | GO:1903320 | regulation of protein modification by small protein conjugation or removal | 15 | 3 | 0.38 | 0.00598 | 0.00641 |
| L: 2x up 4x | GO:1904666 | regulation of ubiquitin protein ligase activity | 15 | 3 | 0.38 | 0.00598 | 0.00641 |
| L: 2x up 4x | GO:0007264 | small GTPase mediated signal transduction | 46 | 5 | 1.17 | 0.00614 | 0.00641 |

Continued Table S9

| Group | GO.ID | Term | Annotated | Significant | Expected | Fisher.*P* | Adjust.*P* |
|---|---|---|---|---|---|---|---|
| L: 2x up 4x | GO:1901564 | **organonitrogen compound metabolic process** | 6333 | 187 | 161.65 | 0.00730 | 0.00730 |
| L: 2x dw 4x | GO:0006952 | **defense response** | 253 | 26 | 10.84 | 0.00004 | 0.00074 |
| L: 2x dw 4x | GO:0000184 | **nuclear-transcribed mRNA catabolic process, nonsense-mediated decay** | 17 | 5 | 0.73 | 0.00057 | 0.00057 |
| L: 2x dw 4x | GO:0009311 | oligosaccharide metabolic process | 84 | 11 | 3.6 | 0.00089 | 0.00268 |
| L: 2x dw 4x | GO:0010215 | cellulose microfibril organization | 28 | 6 | 1.2 | 0.00102 | 0.00268 |
| L: 2x dw 4x | GO:0030198 | extracellular matrix organization | 28 | 6 | 1.2 | 0.00102 | 0.00268 |
| L: 2x dw 4x | GO:0043062 | **extracellular structure organization** | 28 | 6 | 1.2 | 0.00102 | 0.00268 |
| L: 2x dw 4x | GO:0070726 | cell wall assembly | 28 | 6 | 1.2 | 0.00102 | 0.00268 |
| L: 2x dw 4x | GO:0071668 | **plant-type cell wall assembly** | 28 | 6 | 1.2 | 0.00102 | 0.00268 |
| L: 2x dw 4x | GO:0043603 | **cellular amide metabolic process** | 1152 | 71 | 49.35 | 0.00123 | 0.00287 |
| L: 2x dw 4x | GO:0009832 | plant-type cell wall biogenesis | 31 | 6 | 1.33 | 0.00178 | 0.00374 |
| L: 2x dw 4x | GO:0006518 | peptide metabolic process | 1103 | 67 | 47.25 | 0.00240 | 0.00458 |
| L: 2x dw 4x | GO:0009245 | lipid A biosynthetic process | 8 | 3 | 0.34 | 0.00373 | 0.00522 |
| L: 2x dw 4x | GO:0046493 | lipid A metabolic process | 8 | 3 | 0.34 | 0.00373 | 0.00522 |
| L: 2x dw 4x | GO:1901269 | lipooligosaccharide metabolic process | 8 | 3 | 0.34 | 0.00373 | 0.00522 |
| L: 2x dw 4x | GO:1901271 | lipooligosaccharide biosynthetic process | 8 | 3 | 0.34 | 0.00373 | 0.00522 |
| L: 2x dw 4x | GO:0019538 | protein metabolic process | 5121 | 252 | 219.38 | 0.00474 | 0.00622 |
| L: 2x dw 4x | GO:0009312 | oligosaccharide biosynthetic process | 64 | 8 | 2.74 | 0.00581 | 0.00715 |
| L: 2x dw 4x | GO:0071669 | plant-type cell wall organization or biogenesis | 107 | 11 | 4.58 | 0.00613 | 0.00715 |
| L: 2x dw 4x | GO:0006749 | glutathione metabolic process | 125 | 12 | 5.35 | 0.00741 | 0.00819 |
| L: 2x dw 4x | GO:0000956 | nuclear-transcribed mRNA catabolic process | 43 | 6 | 1.84 | 0.00956 | 0.00956 |
| L: 2x dw 4x | GO:0005991 | trehalose metabolic process | 43 | 6 | 1.84 | 0.00956 | 0.00956 |
| R: 2x up 4x | GO:0009065 | glutamine family amino acid catabolic process | 7 | 4 | 0.21 | 0.00003 | 0.00120 |
| R: 2x up 4x | GO:0007165 | **signal transduction** | 666 | 38 | 19.72 | 0.00009 | 0.00150 |

Continued Table S9

| Group | GO.ID | Term | Annotated | Significant | Expected | Fisher.*P* | Adjust.*P* |
|---|---|---|---|---|---|---|---|
| R: 2x up 4x | GO:0023052 | signaling | 667 | 38 | 19.74 | 0.00009 | 0.00150 |
| R: 2x up 4x | GO:0007154 | **cell communication** | 679 | 38 | 20.1 | 0.00014 | 0.00168 |
| R: 2x up 4x | GO:0009694 | **jasmonic acid metabolic process** | 5 | 3 | 0.15 | 0.00025 | 0.00200 |
| R: 2x up 4x | GO:0009695 | jasmonic acid biosynthetic process | 5 | 3 | 0.15 | 0.00025 | 0.00200 |
| R: 2x up 4x | GO:0009064 | glutamine family amino acid metabolic process | 61 | 8 | 1.81 | 0.00041 | 0.00281 |
| R: 2x up 4x | GO:0044281 | small molecule metabolic process | 1421 | 64 | 42.06 | 0.00049 | 0.00294 |
| R: 2x up 4x | GO:0050896 | **response to stimulus** | 1826 | 78 | 54.05 | 0.00057 | 0.00302 |
| R: 2x up 4x | GO:0006793 | phosphorus metabolic process | 2713 | 108 | 80.31 | 0.00063 | 0.00302 |
| R: 2x up 4x | GO:0051716 | cellular response to stimulus | 1113 | 52 | 32.95 | 0.00075 | 0.00327 |
| R: 2x up 4x | GO:0006796 | phosphate-containing compound metabolic process | 2707 | 107 | 80.13 | 0.00086 | 0.00344 |
| R: 2x up 4x | GO:0009640 | photomorphogenesis | 17 | 4 | 0.5 | 0.00133 | 0.00480 |
| R: 2x up 4x | GO:0016310 | phosphorylation | 2032 | 83 | 60.15 | 0.00143 | 0.00480 |
| R: 2x up 4x | GO:0006468 | protein phosphorylation | 1889 | 78 | 55.92 | 0.00150 | 0.00480 |
| R: 2x up 4x | GO:0031124 | mRNA 3'-end processing | 20 | 4 | 0.59 | 0.00252 | 0.00654 |
| R: 2x up 4x | GO:0006562 | proline catabolic process | 3 | 2 | 0.09 | 0.00257 | 0.00654 |
| R: 2x up 4x | GO:0044283 | small molecule biosynthetic process | 463 | 25 | 13.71 | 0.00297 | 0.00654 |
| R: 2x up 4x | GO:0008037 | cell recognition | 67 | 7 | 1.98 | 0.00359 | 0.00654 |
| R: 2x up 4x | GO:0048544 | recognition of pollen | 67 | 7 | 1.98 | 0.00359 | 0.00654 |
| R: 2x up 4x | GO:0019752 | **carboxylic acid metabolic process** | 871 | 40 | 25.78 | 0.00404 | 0.00654 |
| R: 2x up 4x | GO:0043436 | oxoacid metabolic process | 873 | 40 | 25.84 | 0.00420 | 0.00654 |
| R: 2x up 4x | GO:0006082 | organic acid metabolic process | 877 | 40 | 25.96 | 0.00453 | 0.00654 |
| R: 2x up 4x | GO:0046394 | carboxylic acid biosynthetic process | 330 | 19 | 9.77 | 0.00460 | 0.00654 |
| R: 2x up 4x | GO:0000103 | sulfate assimilation | 12 | 3 | 0.36 | 0.00465 | 0.00654 |
| R: 2x up 4x | GO:0006378 | mRNA polyadenylation | 12 | 3 | 0.36 | 0.00465 | 0.00654 |
| R: 2x up 4x | GO:0031398 | positive regulation of protein ubiquitination | 12 | 3 | 0.36 | 0.00465 | 0.00654 |

Continued Table S9

| Group | GO.ID | Term | Annotated | Significant | Expected | Fisher.*P* | Adjust.*P* |
|---|---|---|---|---|---|---|---|
| R: 2x up 4x | GO:0051443 | positive regulation of ubiquitin-protein transferase activity | 12 | 3 | 0.36 | 0.00465 | 0.00654 |
| R: 2x up 4x | GO:1903322 | positive regulation of protein modification by small protein conjugation or removal | 12 | 3 | 0.36 | 0.00465 | 0.00654 |
| R: 2x up 4x | GO:1904668 | positive regulation of ubiquitin protein ligase activity | 12 | 3 | 0.36 | 0.00465 | 0.00654 |
| R: 2x up 4x | GO:0006527 | arginine catabolic process | 4 | 2 | 0.12 | 0.00504 | 0.00654 |
| R: 2x up 4x | GO:0031440 | regulation of mRNA 3'-end processing | 4 | 2 | 0.12 | 0.00504 | 0.00654 |
| R: 2x up 4x | GO:0031441 | negative regulation of mRNA 3'-end processing | 4 | 2 | 0.12 | 0.00504 | 0.00654 |
| R: 2x up 4x | GO:0050686 | negative regulation of mRNA processing | 4 | 2 | 0.12 | 0.00504 | 0.00654 |
| R: 2x up 4x | GO:1900363 | regulation of mRNA polyadenylation | 4 | 2 | 0.12 | 0.00504 | 0.00654 |
| R: 2x up 4x | GO:1900364 | negative regulation of mRNA polyadenylation | 4 | 2 | 0.12 | 0.00504 | 0.00654 |
| R: 2x up 4x | GO:1903312 | negative regulation of mRNA metabolic process | 4 | 2 | 0.12 | 0.00504 | 0.00654 |
| R: 2x up 4x | GO:0006520 | cellular amino acid metabolic process | 562 | 28 | 16.64 | 0.00527 | 0.00666 |
| R: 2x up 4x | GO:1901605 | alpha-amino acid metabolic process | 267 | 16 | 7.9 | 0.00615 | 0.00749 |
| R: 2x up 4x | GO:1901606 | alpha-amino acid catabolic process | 40 | 5 | 1.18 | 0.00624 | 0.00749 |
| R: 2x up 4x | GO:0016053 | organic acid biosynthetic process | 369 | 20 | 10.92 | 0.00713 | 0.00835 |
| R: 2x up 4x | GO:0006020 | inositol metabolic process | 14 | 3 | 0.41 | 0.00736 | 0.00841 |
| R: 2x up 4x | GO:0009063 | cellular amino acid catabolic process | 42 | 5 | 1.24 | 0.00769 | 0.00858 |
| R: 2x up 4x | GO:0032502 | developmental process | 276 | 16 | 8.17 | 0.00835 | 0.00900 |
| R: 2x up 4x | GO:0031396 | regulation of protein ubiquitination | 15 | 3 | 0.44 | 0.00900 | 0.00900 |
| R: 2x up 4x | GO:0051438 | regulation of ubiquitin-protein transferase activity | 15 | 3 | 0.44 | 0.00900 | 0.00900 |
| R: 2x up 4x | GO:1903320 | regulation of protein modification by small protein conjugation or removal | 15 | 3 | 0.44 | 0.00900 | 0.00900 |
| R: 2x up 4x | GO:1904666 | regulation of ubiquitin protein ligase activity | 15 | 3 | 0.44 | 0.00900 | 0.00900 |
| R: 2x dw 4x | GO:0010468 | **regulation of gene expression** | 1895 | 110 | 72.5 | 0.00000 | 0.00006 |
| R: 2x dw 4x | GO:0007623 | **circadian rhythm** | 29 | 8 | 1.11 | 0.00001 | 0.00006 |
| R: 2x dw 4x | GO:0048511 | rhythmic process | 29 | 8 | 1.11 | 0.00001 | 0.00006 |

Continued Table S9

| Group | GO.ID | Term | Annotated | Significant | Expected | Fisher.*P* | Adjust.*P* |
|---|---|---|---|---|---|---|---|
| R: 2x dw 4x | GO:0060255 | regulation of macromolecule metabolic process | 2010 | 114 | 76.9 | 0.00001 | 0.00006 |
| R: 2x dw 4x | GO:0050789 | **regulation of biological process** | 2888 | 153 | 110.49 | 0.00001 | 0.00006 |
| R: 2x dw 4x | GO:0019222 | regulation of metabolic process | 2035 | 114 | 77.86 | 0.00002 | 0.00008 |
| R: 2x dw 4x | GO:0000184 | nuclear-transcribed mRNA catabolic proces, nonsense-mediated decay | 17 | 6 | 0.65 | 0.00003 | 0.00003 |
| R: 2x dw 4x | GO:0065007 | biological regulation | 3033 | 156 | 116.04 | 0.00004 | 0.00015 |
| R: 2x dw 4x | GO:0019219 | regulation of nucleobase-containing compound metabolic process | 1747 | 96 | 66.84 | 0.00017 | 0.00044 |
| R: 2x dw 4x | GO:0006355 | regulation of DNA-templated transcription | 1703 | 94 | 65.16 | 0.00017 | 0.00044 |
| R: 2x dw 4x | GO:1903506 | **regulation of nucleic acid-templated transcription** | 1703 | 94 | 65.16 | 0.00017 | 0.00044 |
| R: 2x dw 4x | GO:2001141 | regulation of RNA biosynthetic process | 1703 | 94 | 65.16 | 0.00017 | 0.00044 |
| R: 2x dw 4x | GO:0051252 | regulation of RNA metabolic process | 1731 | 95 | 66.23 | 0.00019 | 0.00045 |
| R: 2x dw 4x | GO:0050794 | regulation of cellular process | 2633 | 134 | 100.74 | 0.00025 | 0.00053 |
| R: 2x dw 4x | GO:0010556 | regulation of macromolecule biosynthetic process | 1745 | 95 | 66.76 | 0.00026 | 0.00053 |
| R: 2x dw 4x | GO:0009889 | regulation of biosynthetic process | 1751 | 95 | 66.99 | 0.00029 | 0.00053 |
| R: 2x dw 4x | GO:0031326 | regulation of cellular biosynthetic process | 1751 | 95 | 66.99 | 0.00029 | 0.00053 |
| R: 2x dw 4x | GO:0010467 | gene expression | 3701 | 178 | 141.6 | 0.00038 | 0.00065 |
| R: 2x dw 4x | GO:0051171 | **regulation of nitrogen compound metabolic process** | 1862 | 99 | 71.24 | 0.00043 | 0.00070 |
| R: 2x dw 4x | GO:0031323 | regulation of cellular metabolic process | 1865 | 99 | 71.35 | 0.00046 | 0.00071 |
| R: 2x dw 4x | GO:0080090 | regulation of primary metabolic process | 1872 | 99 | 71.62 | 0.00052 | 0.00077 |
| R: 2x dw 4x | GO:0006952 | defense response | 253 | 21 | 9.68 | 0.00076 | 0.00107 |
| R: 2x dw 4x | GO:0009059 | macromolecule biosynthetic process | 3249 | 155 | 124.31 | 0.00145 | 0.00195 |
| R: 2x dw 4x | GO:0006351 | DNA-templated transcription | 1947 | 99 | 74.49 | 0.00187 | 0.00232 |
| R: 2x dw 4x | GO:0097659 | nucleic acid-templated transcription | 1947 | 99 | 74.49 | 0.00187 | 0.00232 |
| R: 2x dw 4x | GO:0032774 | RNA biosynthetic process | 1951 | 99 | 74.64 | 0.00199 | 0.00237 |
| R: 2x dw 4x | GO:0016070 | RNA metabolic process | 2802 | 133 | 107.2 | 0.00405 | 0.00465 |

Continued Table S9

| Group | GO.ID | Term | Annotated | Significant | Expected | Fisher.*P* | Adjust.*P* |
|---|---|---|---|---|---|---|---|
| R: 2x dw 4x | GO:0000956 | nuclear-transcribed mRNA catabolic process | 43 | 6 | 1.65 | 0.00560 | 0.00620 |
| R: 2x dw 4x | GO:0050896 | **response to stimulus** | 1826 | 90 | 69.86 | 0.00708 | 0.00757 |
| R: 2x dw 4x | GO:0009416 | **response to light stimulus** | 77 | 8 | 2.95 | 0.00909 | 0.00939 |
| R: 2x dw 4x | GO:0044249 | cellular biosynthetic process | 4303 | 191 | 164.63 | 0.00995 | 0.00995 |
